# *MECP2* duplication and mutations impair NSCs differentiation via miR-197 regulated ADAM10

**DOI:** 10.1101/312983

**Authors:** Yu-Meng Wang, Yu-Fang Zheng, Si-Yu Yang, Zhang-Min Yang, Lin-Na Zhang, Yan-Qin He, Xiao-Hong Gong, Dong Liu, Richard H. Finnell, Zi-Long Qiu, Ya-Song Du, Hong-Yan Wang

## Abstract

How *MECP2* (Methyl-CpG-binding protein 2) duplication affects cortex development remains elusive. We found that elevated MeCP2 expression promotes neurogenesis during cortex development in Tg(*MECP2*) mouse brain. Ectopic expression of MeCP2 in NPCs inhibits ADAM10 and hence compromises the NOTCH pathway during NPC differentiation. MeCP2 up-regulates miR-197 to down-regulate ADAM10. The enhanced NPC differentiation/migration in Tg(MECP2) embryonic brain can be repressed by overexpression of ADAM10 or a miR-197 inhibitor.

Consistently, the reduced neurogenesis induced by three rare MeCP2 missense mutations (H371R, E394K, G428S) identified in a Han Chinese autism spectrum disorders (ASD) cohort, can be reversed by miR-197 both in vitro and in vivo. Our results revealed that a regulatory axis involving MeCP2, miR-197, ADAM10, and NOTCH signaling is critical for neurogenesis, which is affected by both MeCP2 duplication and mutation.

## Introduction

*MECP2* (methyl-CpG binding protein 2), an X-linked gene encoding the methyl-cytosine binding protein MeCP2, is associated with two severe neurological disorders, Rett syndrome (RTT) and *MECP2* duplication syndrome (MDS), which result from loss and gain of function of *MECP2*, respectively. Mutations in *MECP2* were first identified as the monogenic cause for RTT (1). Although clinically distinguished from Autism Spectrum Disorder (ASD) by the Diagnostic and Statistical Manual of Mental Disorders (DSM-5) (2), autistic features are often observed (>60%) in RTT patients (3, 4). The *Mecp2* knockout mice (*Mecp2*-/y) also showed excellent replication/representation of RTT and ASD phenotypes (5, 6). Recently, MDS, a severe male intellectual disability syndrome, caused by *MECP2* duplication was identified (7, 8), which has ~100% penetrance of autistic-like behaviors (9, 10). Transgenic mouse models for MDS (11, 12) and lentivirus-based transgenic monkey (13) both showed severe autistic-like and compulsive repetitive behaviors. Therefore, it has been suggested that the functional and dosage variations of MeCP2 are tightly associated with autistic features (4, 14). Exploring the functional conversions of *MECP2* duplications and mutations may provide useful insight into the mechanisms underlying the development of ASD phenotypes (4, 14).

MeCP2 was originally identified as a transcriptional repressor (15, 16), but recent studies have shown it has diverse functions, including transcription activation (17), mRNA splicing (18, 19), and microRNA (miRNA) processing (20, 21), to name but a few. As both RTT and ASD patients show symptoms shortly after birth, most functional studies of MeCP2 have been focused on postnatal events such as dendritic arborization (22-24), synapse formation and plasticity (24-26), and adult neurogenesis (27-29). However, the function of MeCP2 during embryonic CNS development is still elusive.

MeCP2 is widely and highly expressed in the developing central nervous system (CNS), including both the early neural tube and embryonic forebrain, in zebrafish, chicken, and mouse (30-33). However, embryonic development seems to be unaffected in *Mecp2* knockout mice (6, 34). But study in *Xenopus* showed that MeCP2 promoted neurogenesis of *Xenopus* embryos, while the RTT mutant R168X failed to do so (35). A recent study in the monkey showed that Talent-edited mutation of *MECP2* caused embryonic lethality in male mutant monkeys (36). Furthermore, human iPSCs generated from RTT patients with dysfunctional MeCP2 have abnormal neurogenesis and gliogenesis (37, 38). Therefore, it is likely that MeCP2 has an important role during early CNS development.

In the present study, we demonstrated that overexpressed MeCP2 promoted neurogenesis in the embryonic brain in the MDS model Tg(*MECP2*) mouse. Our results revealed a novel mechanism involving miR-197, ADAM10 (A disintergrin and metalloprotease 10), and NOTCH signaling as a critical regulatory axis for the enhanced neurogenesis induced by MeCP2. Furthermore, we identified three rare missense *MECP2* mutations (H371R, E394K, G428S) in ASD patients in our Han Chinese ASD cohort, which are novel in the East Asian population according to ExAC (39). These *MECP2* mutations resulted in dysfunctional regulation of miR-197 and neurogenesis. Surprisingly, while the inhibitor of miR-197 could reverse the effect of overexpressed MeCP2, the overexpression of miR-197 could reverse the NPCs differentiation defects caused by *MECP2* mutations both *in vitro* and *in vivo*. Our results revealed a novel regulatory pathway via miR-197 by which MeCP2 acting on ADAM10/NOTCH signaling, implicating that molecules in this pathway are important for the etiology of MDS and possibly for ASD.

## Results

### Neurogenesis is enhanced in Tg(*MECP2*) mouse fetal brain

To determine the effect of *MECP2* duplication on cell fate *in vivo* in fetal brain, we investigated a Tg(*MECP2*) mouse line which had previously been used as a model for MDS (11), as it contains an extra copy of human *MECP2* and exhibits approximately doubled MeCP2 levels in the brain (Fig. 2D). During cortical neurogenesis in the mouse, NSCs/NPCs in the ventricular (VZ) and subventricular (SVZ) zones are differentiated into immature neurons and migrate radially to the cortical plate (CP) to become mature neurons (40, 41). For quantification of cell fate in wild-type (WT) and Tg(*MECP2*) cortex, immunofluorescent staining on E18.5 and P7 brain sections with different markers were performed. The results revealed that there are significantly more Satb2^+^ (a cortical neuron marker) cells in CP layer and significantly less Sox2^+^ and Tbr2^+^ (progenitor markers) cells in VZ/SVZ layer of E18.5 Tg*(MECP2)* mice compared to WT littermates (Fig. 1A). By P7, there are still significantly more Satb2^+^ and Tbr1^+^ neurons present in P7 Tg*(MECP2)* mice cortex than WT cortex (Fig. 1B).

**Figure 1.**
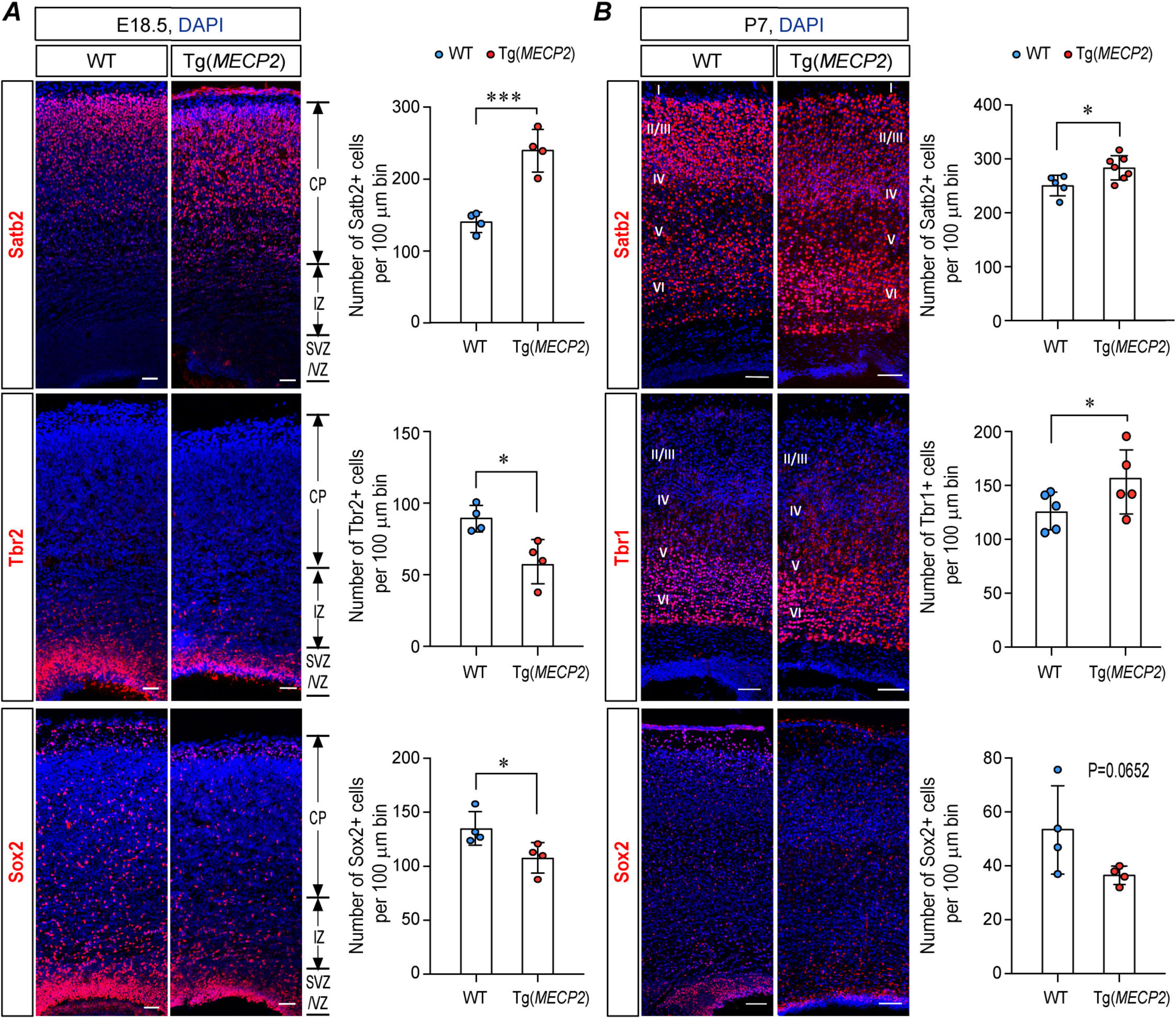
*MECP2* duplication promotes neurogenesis in Tg*(MECP2)* FVB mice cortex. Coronal brain sections from E18.5 **(A)** and P7 **(B)** Tg(*MECP2*) FVB mice or WT littermates were stained with cortical neuron marker Satb2, deep layer neuron marker Tbr1, and progenitor markers Sox2 and Tbr2. N≥ 4. DAPI (blue) was used for nuclear staining. The numbers of positive cells within 100μm bin were counted. Representative sections are shown in left and statistical analysis are shown in right panels, respectively. All statistic data represent means ± SEM. * *p*<0.05, *** *p*<0.001. Scale bar is 50μm in E18.5 and 100μm in P7.

**Figure 2.**
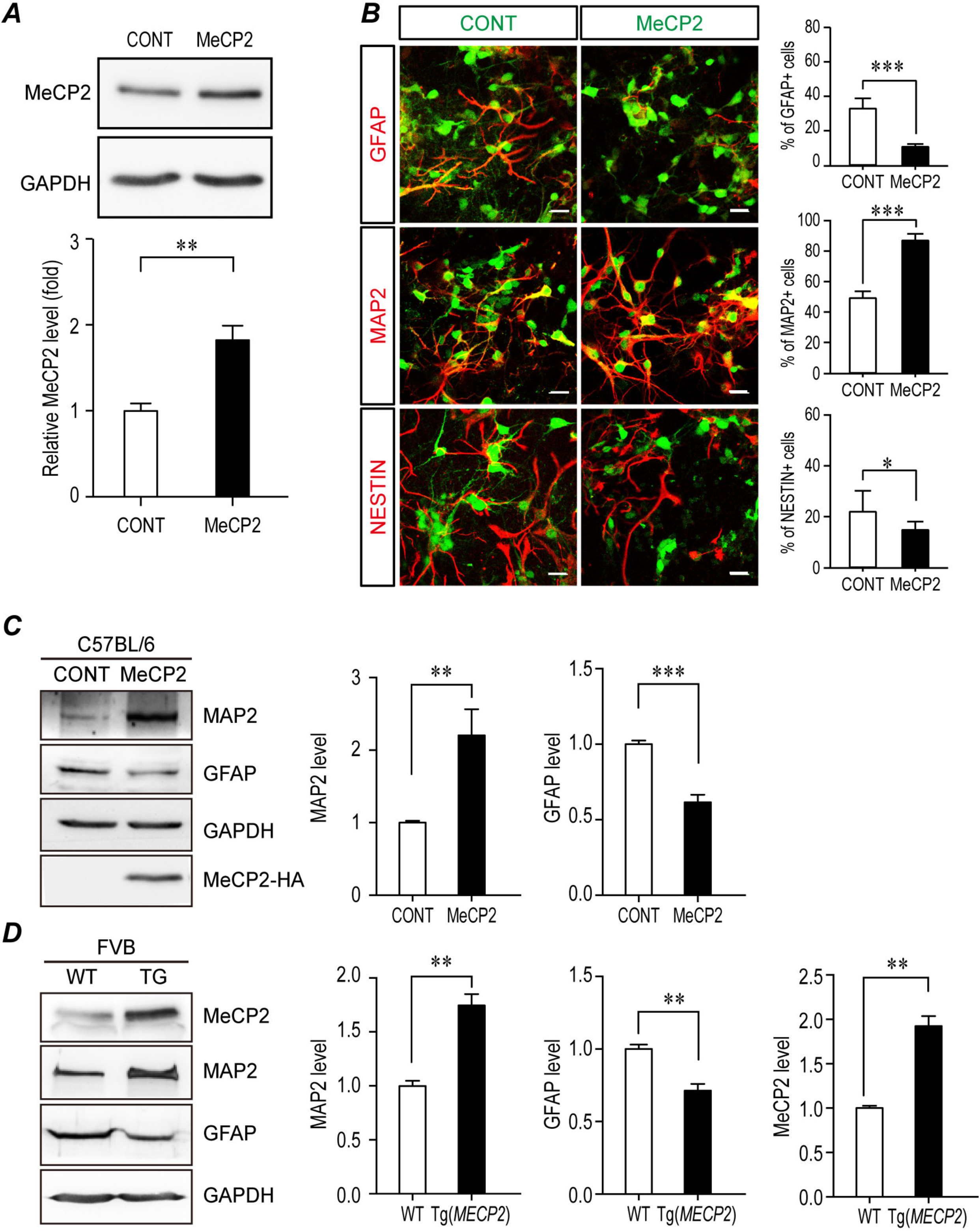
*MECP2* duplication promotes neurogenesis of cultured NPCs. Mouse primary NPCs isolated from C57BL/6 mouse E12.5 embryonic cortex were infected with MeCP2 expressing lentivirus and cultured for 72hrs **(A-C)**. Cell lysates were subjected to western blotting or immunofluorescent staining. **(A)** Exogenous expressing of MeCP2 doubled its expression level in NPCs. GAPDH was used as loading control. N=3. Representative blot is shown on the top panel and statistical analysis for MeCP2 levels are shown in the bottom panel. **(B)** Immunofluorescent staining for GFAP, MAP2, and NESTIN were performed on NPCs. N=9. The MeCP2 and control viruses contain EGFP and showed here as green. The staining for MAP2, GFAP, and NESTIN are labeled with pseudo-colored red. Representative images are shown on the left and the percentage of Nestin^+^, MAP2^+^ and GFAP^+^ cells within EGFP^+^ cells are shown on the right panels respectively. Scale bar is 50μm. **(C)** Western blot analysis for MAP2, GFAP, and HA labeled MeCP2. GAPDH was used as loading control. N=9. Representative blot is shown on the left, and statistical analysis for MAP2 and GFAP levels are shown in the right panels, respectively. **(D)** Mouse primary NPCs isolated from FVB WT and Tg(*MECP2*) mouse E12.5 embryonic cortex were directly cultured for 72hrs. Cell lysates were subjected to western blot analysis for both MeCP2 and differentiation markers, MAP2 and GFAP. GAPDH was used as loading control. N=3. Representative blot is shown in left and statistical analysis for MeCP2, MAP2 and GFAP levels are shown in right panels, respectively. All statistic data represent means ± SEM. * *p*<0.05, ** *p*<0.01, *** *p*<0.001.

Primary NPCs were isolated from E12.5 mouse cortex to further examine the *in vitro* effect of MeCP2 overexpression on NPCs differentiation. WT NPCs from C57BL/6 mice were infected with MeCP2 lentivirus after seeding on the culture dish. The exogenous expression doubled the expression levels of MeCP2 in those NPCs (Fig. 2A). Both transiently overexpressed MeCP2 in C57BL/6 NPCs and elevated MeCP2 in transgenic Tg(*MECP2*) mice significantly up-regulated the level of the neuronal marker MAP2, and down-regulated the level of the glia marker GFAP in NPCs (Fig. 2C&D). Immunofluorescent staining on cultured NPCs infected by lentivirus also showed significantly more MAP2^+^ and less GFAP^+^ cells with MeCP2 overexpression (Fig. 2B), which was consistent with previous reports (42). Taken together, the results demonstrate that elevated MeCP2 expression affects NPCs cell fate and promotes neurogenesis.

### ADAM10 is downstream of MeCP2 in the control of neurogenesis

Since NOTCH is a key molecule for NSCs fate (43, 44), we next examined the levels of several key molecules in the NOTCH pathway in primary cultured NPCs after overexpressing MeCP2. The levels of two NOTCH ligands JAG1 and DLL1, the full-length receptor NOTCH1, the activated NOTCH intracellular domain (NICD), and the two rate-limiting S2 enzymes for NOTCH cleavage, ADAM10 and ADAM17, were examined by Western blot analysis in primary NPCs transfected with either MeCP2 or an empty control vector. The results showed that the levels of NICD and ADAM10 were significantly down-regulated by MeCP2 expression in NPCs (Fig. 3A), while ADAM17, full-length NOTCH1, JAG1, and DLL1 were not significantly affected (Fig. S1A-E). These results suggest that MeCP2 might inhibit NOTCH signaling through the down-regulation of ADAM10, in order to regulate NPC development.

**Figure 3.**
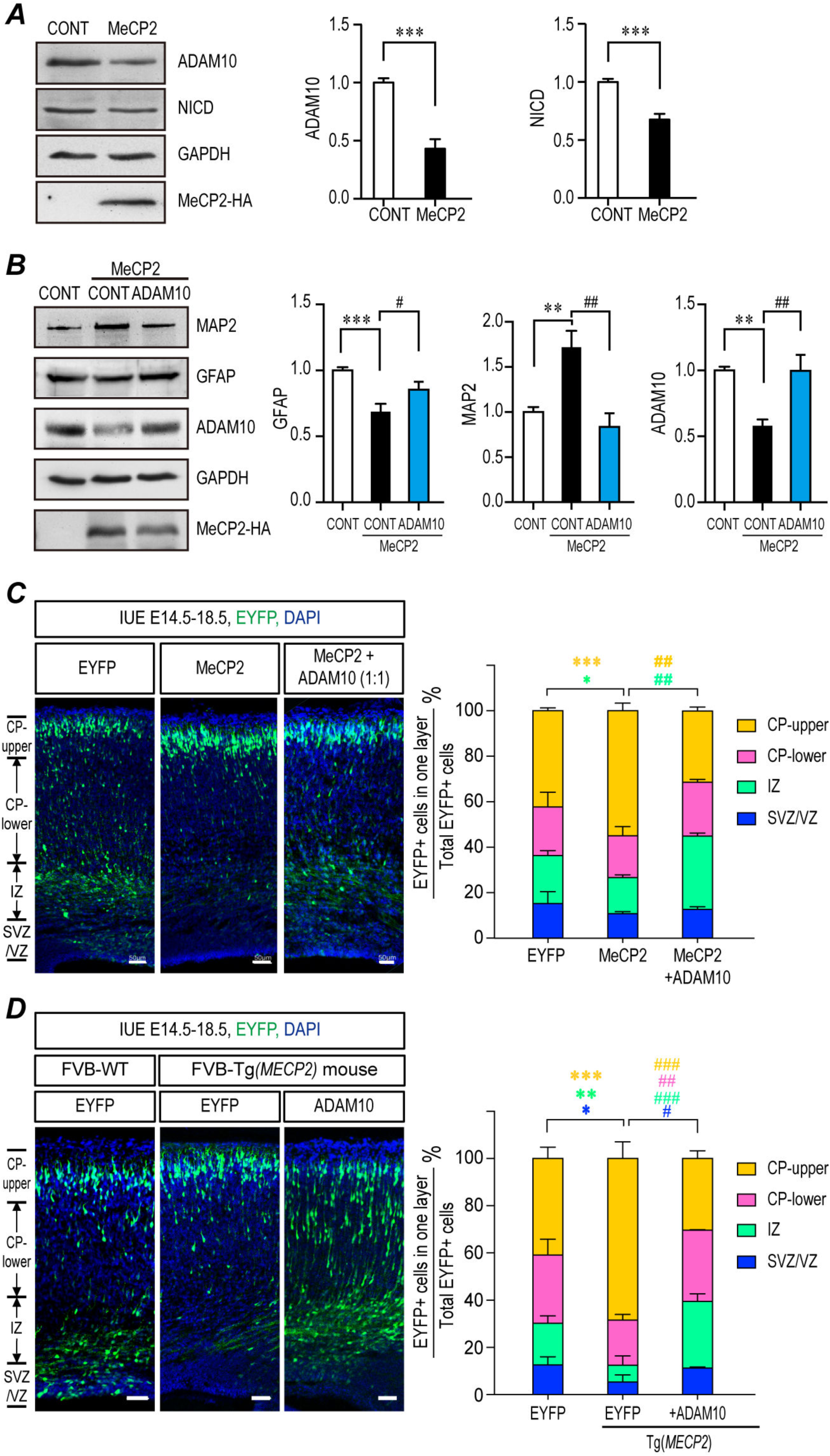
ADAM10 is a critical down-stream molecule of MeCP2 in regulating NPCs differentiation. **(A)** Mouse primary NPCs isolated from E12.5 C57BL/6 mouse embryonic cortex were transfected with either empty vector or MeCP2 expressing plasmids and cultured for 72hrs. Cell lysates were subjected to Western blot analysis for ADAM10 and NICD. GAPDH was used as a loading control. N≥5. Representative blots are shown on the left and statistical analyses for ADAM10 and NICD were presented on the right panels. **(B)** Overexpression of ADAM10 together with WT MeCP2 could reverse the differentiation effects of MeCP2 in cultured NPCs. Mouse primary NPCs isolated from E12.5 C57BL/6 mouse embryonic cortex were transfected with HA tagged MeCP2 expressing plasmids with either control vector or ADAM10 expressing plasmid. 72hrs later, NPCs lysates were subjected to Western blot analysis for MAP2, GFAP, ADAM10, and HA. GAPDH was used as a loading control. N≥5. Representative blots are shown on the left panel and statistical analyses for GFAP, MAP2, and ADAM10 were presented on the right panel, respectively. Overexpression of ADAM10 could reverse the differentiation effects of MeCP2 in electroporated fetal brains **(C-D)**. **(C)** C57BL/6 mouse were electroporated at E14.5 with either MeCP2 or MeCP2 with ADAM10 at 1:1 ratio. All samples were collected at E18.5 for sectioning and immunostaining. Each condition was repeated in four different embryos from four pregnant female mice (N=4). Representative brain sections are presented on the left and the EYFP^+^ cells in each layer were counted and compared to the total EYFP^+^ cells. The statistical analyses for cells in each layer are shown on the right panel. **(D)** *Tg(MECP2)* or WT FVB mice cortex were electroporated with EYFP or ADAM10 with EYFP plasmid (N=4). Representative brain sections are presented on the left and the EYFP^+^ cells in each layer were counted and compared to the total EYFP^+^ cells. The statistical analyses for cells in each layer are presented on the right panel. DAPI (blue) was used for nuclear staining. Scale bar is 50μm in E18.5 and 100μm in P7. All statistic data represent means ± SEM. *, # *p*<0.05, **, ## *p*<0.01, ***, ### *p*<0.001.

To study the relationship between ADAM10 and MeCP2, we expressed ADAM10 together with MeCP2 either in primary cultured NPCs or in embryonic brain. Expression of ADAM10 together with MeCP2 in NPCs reversed the effect of MeCP2 as indicated by the restoration of MAP2 and GFAP levels close to those of controls (Fig. 3B). The *in vivo* effect of ADAM10 on MeCP2 promoted neurogenesis was also tested in the developing cortex by *in utero* electroporation (IUE) experiments. Mouse embryonic brain were electroporated at embryonic day 14.5 (E14.5) and inspected at E18.5. There were significantly more EYFP^+^ cells in CP layer in the Tg*(MECP2)* cortex (Fig. 3D) as well as in C57BL/6 WT cortex ectopically overexpressing MeCP2 (Fig. 3C). Further, immunofluorescent staining showed that those EYFP^+^ cells migrated into the CP layer were positive for the neuronal markers NeuN and Tuj1; whereas those cells remaining in the VZ/SVZ layer were positive for the progenitor markers Sox2 and Tbr2 (Fig. S2). When ADAM10 was co-expressed with MeCP2, it could reverse the effect of MeCP2 on neurogenesis, resulting in significantly fewer cells reaching the upper CP layer, compared to MeCP2 overexpression cortex (Fig. 3C&D). Such effects were not observed when ADAM17 was co-expressed with MeCP2 (Fig. S1F&G). These results suggest that ADAM10 is functionally downstream of MeCP2 during neurogenesis.

### MeCP2 down-regulates ADAM10 expression through up-regulating miR-197

We next investigated how MeCP2 regulates ADAM10. Initially, we confirmed the regulation of ADAM10 by MeCP2 in human cells. Similar to cultured mouse NPCs, the ADAM10 protein level was also significantly down-regulated (~53%) by MeCP2 overexpression in a human glioblastoma cell line U251 (Fig. 4A). The mRNA level of ADAM10 was down-regulated ~19% by MeCP2 (Fig. 4B). Since ADAM10 was not identified as a transcriptional target of MeCP2 (17), we screened the potential microRNAs (miRNAs) that might be involved in ADAM10 regulation by MeCP2. Seven miRNAs, identified as the most significantly regulated miRNAs by MeCP2 in the adult NSCs of the *Mecp2*-/y mouse (45), were used for our studies. When inhibitors for each miRNA (e.g. i-197 as an inhibitor of miR-197) were transfected into U251 cells, ADAM10 protein levels were significantly up-regulated by i-222, i-193 and i-197; while the other four inhibitors had no statistically significant effect (Fig. S3A). However, when MeCP2 was co-transfected with miRNA inhibitors, only i-197 and i-193 were able to reverse the MeCP2-induced down-regulation of ADAM10 protein (Fig. 4D & Fig. S3B). The expression profile from miRNAMap shows that hsa-miR-193 is primarily expressed in the muscle, and hsa-miR-197 is highly expressed in the brain (http://mirnamap.mbc.nctu.edu.tw) (46); therefore, miR-197 is the most likely target of MeCP2 in the regulation of ADAM10 expression in the brain.

**Figure 4.**
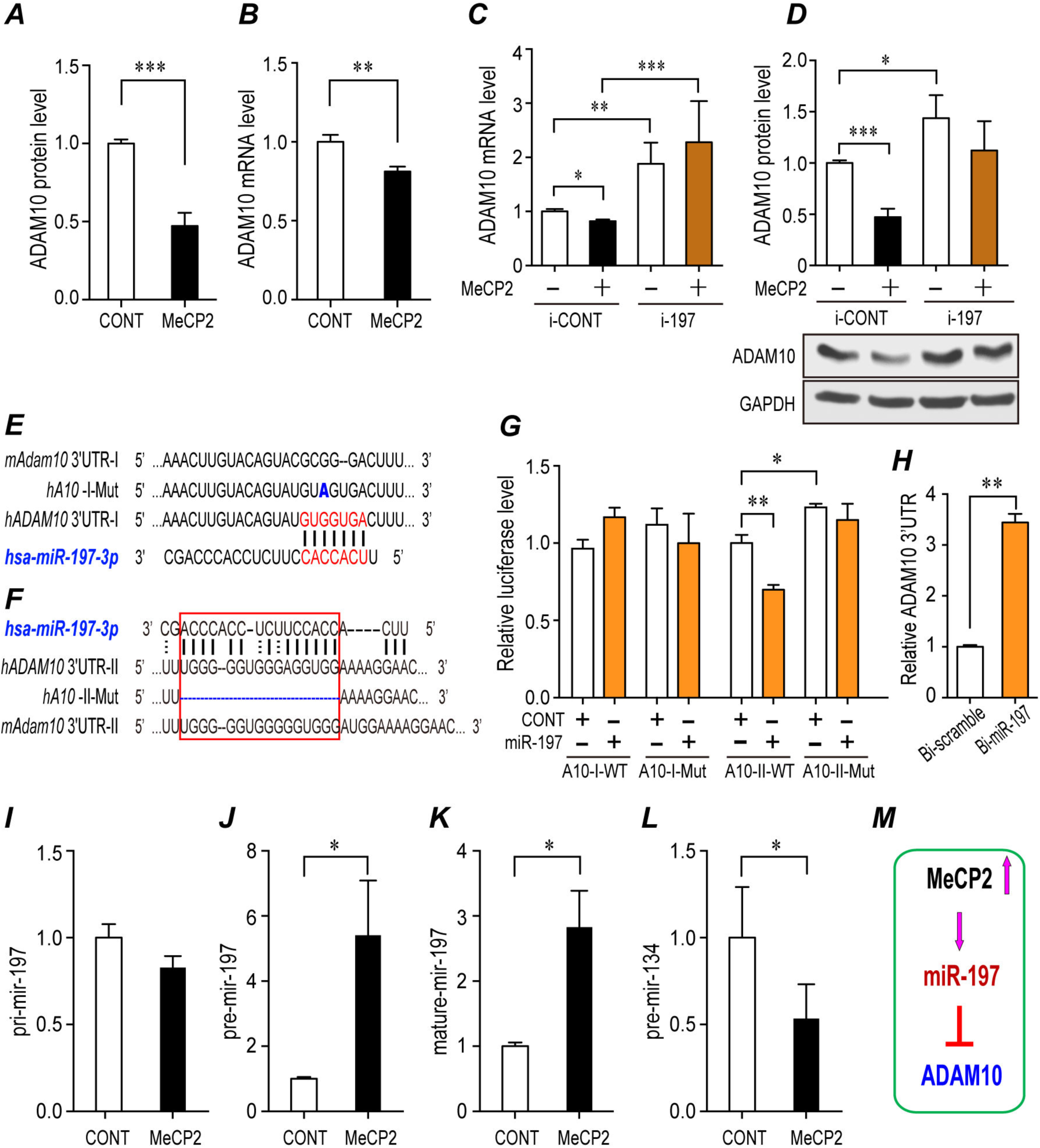
MiR-197 is up-regulated by MeCP2 to down-regulate ADAM10. **(A-B)** Human glioblastoma cells U251 were transfected with empty control or plasmid to overexpress MeCP2, and subjected to either western blotting or qRT-PCR to detect ADAM10 protein **(A)** or mRNA **(B)**. N≥5. **(C-D)** U251 cells were transfected with MeCP2 and a miR-197 inhibitor or scramble control, and subjected to qRT-PCR for ADAM10 mRNA **(C)** or Western blotting for ADAM10 protein **(D)**. N≥3. GAPDH was used as loading control in both experiments. **(E)** A 7mer-m8 miR-197 binding site predicted by TargetscanHuman. The alignment of miR-197 to human and mouse *ADAM10* 3’-UTR and the point mutation G>A in A10-I-Mut are illustrated. **(F)** A miR-197 binding site predicted by RNAhybrid is highly conserved between human and mouse. The deletion mutation in A10-II-Mut was also illustrated. **(G)** Luciferase reporter plasmids for *ADAM10* 3’-UTR were transfected into U251 cells with miR-197 mimics or control. N≥8. **(H)** Biotin-labeled miR-197 and control scramble were transfected into U251 cells and the *ADAM10* 3’-UTR pulled down by miR-197 or control scramble were quantified by qRT-PCR. N=4. **(I-L)** U251 cells were transfected with either control empty vector or MeCP2 expressing plasmids and cultured for 24hrs. RNA from these cells were extracted and subjected for qRT-PCR. N=9. The levels of pri-miR-197, pre-miR-197, mature miR-197, and pre-miR-134 were shown in **(I-L)**, respectively. All statistical data are represented as means ± SEM. * *p*<0.05, ** *p*<0.01, *** *p*<0.001. **(M)** A scheme illustration for MeCP2 regulation on miR-197 and ADAM10.

We went on to investigate the potential target site on *ADAM10* 3’-UTR for miR-197. A 7mer-m8 miR-197 binding site at position 1568-1574 of human *ADAM10* 3’-UTR was predicted by TargetscanHuman (http://www.targetscan.org/vert_71/) (47), but it is poorly conserved between human and mouse (position 1563-1568 of mouse *Adam10* 3’-UTR) (Fig. 4E). A luciferase reporter A10-I-WT was constructed with a 305bp fragment of human *ADAM10* 3’-UTR covering this predicted position 1568-1574. The G to A point mutation in the seed sequence was also constructed as A10-I-Mut (Fig. 4E). Both constructs were transfected into U251 cells with either miR-197 mimics or mimic control. Surprisingly, the results showed that miR-197 had no effect on either A10-I-WT nor A10-I-Mut (Fig. 4G), rather the point mutation blocked the interaction of human *ADAM10* 3’-UTR with miR-224, which has an overlapped seed sequence at position 1571-1577 (Fig. S4A&B).

By using RNAhybrid analysis (48), a different miR-197 binding site at position 186-201 of human *ADAM10* 3’-UTR was predicted based on the free energy of the miRNA-target-duplex, which is highly conserved between human and mouse (position 185-199 of mouse *Adam10* 3’-UTR) (Fig. 4F). This binding site is different from traditional miRNAs and it was predicted to bind the 3’ side of miR-197. A luciferase reporter A10-II-WT was constructed with a 402bp fragment of human *ADAM10* 3’-UTR covering positions 186-201. A deletion mutant reporter A10-II-Mut, deleting this untraditional miRNA binding site, was also constructed (Fig. 4F). The relative luciferase level was up-regulated approximately 23% by A10-II-Mut compared to that of A10-II-WT (Fig. 4G). Interestingly, miR-197 mimics significantly down-regulated the relative luciferase level of A10-II-WT but not A10-II-Mut (Fig. 4G). These results suggested a non-canonical interaction of miR-197 to ADAM10 3’-UTR. In support of this, an RNA-IP assay with a biotinylated miR-197 demonstrated that human *ADAM10* 3’-UTR could be pulled-down by miR-197 (Fig. 4H), which confirmed the interaction between miR-197 and *ADAM10* 3’-UTR.

We subsequently sought to determine how MeCP2 regulates miR-197. RT-PCR experiments showed that the levels of pre- and matured miR-197 were significantly up-regulated by approximately 5.4 and 2.7 fold by MeCP2, respectively (Fig. 4J-K), while the level of pri-miR-197 was not affected by MeCP2 (Fig. 3I). This up-regulation of pre-miR-197 by MeCP2 does not affect the down-regulation of pre-miR-134 by MeCP2 (Fig. 4L). Taken together, MeCP2 down-regulated ADAM10 by up-regulating pre-miR-197 (Fig. 4M).

### MiR-197 plays a critical role in MeCP2 mediated neurogenesis

Since MeCP2 expression is capable of up-regulating the expression of miR-197, we overexpressed miR-197 in primary cultured NPCs to better understand its effect on NPCs differentiation. Immunofluorescent staining showed that, like MeCP2, miR-197 overexpression promoted neurogenesis and inhibited gliogenesis, as evidenced by an elevated ratio of MAP2^+^ and decreased ratio of GFAP^+^ cells in the differentiated NPCs (Fig. 5A). The inhibitor of miR-197 (i-197) had opposite effect with decreased ratio of MAP2^+^ and elevated ratio of GFAP^+^ cells (Fig. 5A). Furthermore, the neurogenesis promoting effect of MeCP2 could also be blocked by i-197 when they were co-expressed in NPCs (Fig. 5B). MeCP2 induced down-regulation of ADAM10 was also blocked by i-197 (Fig. S5A).

**Figure 5.**
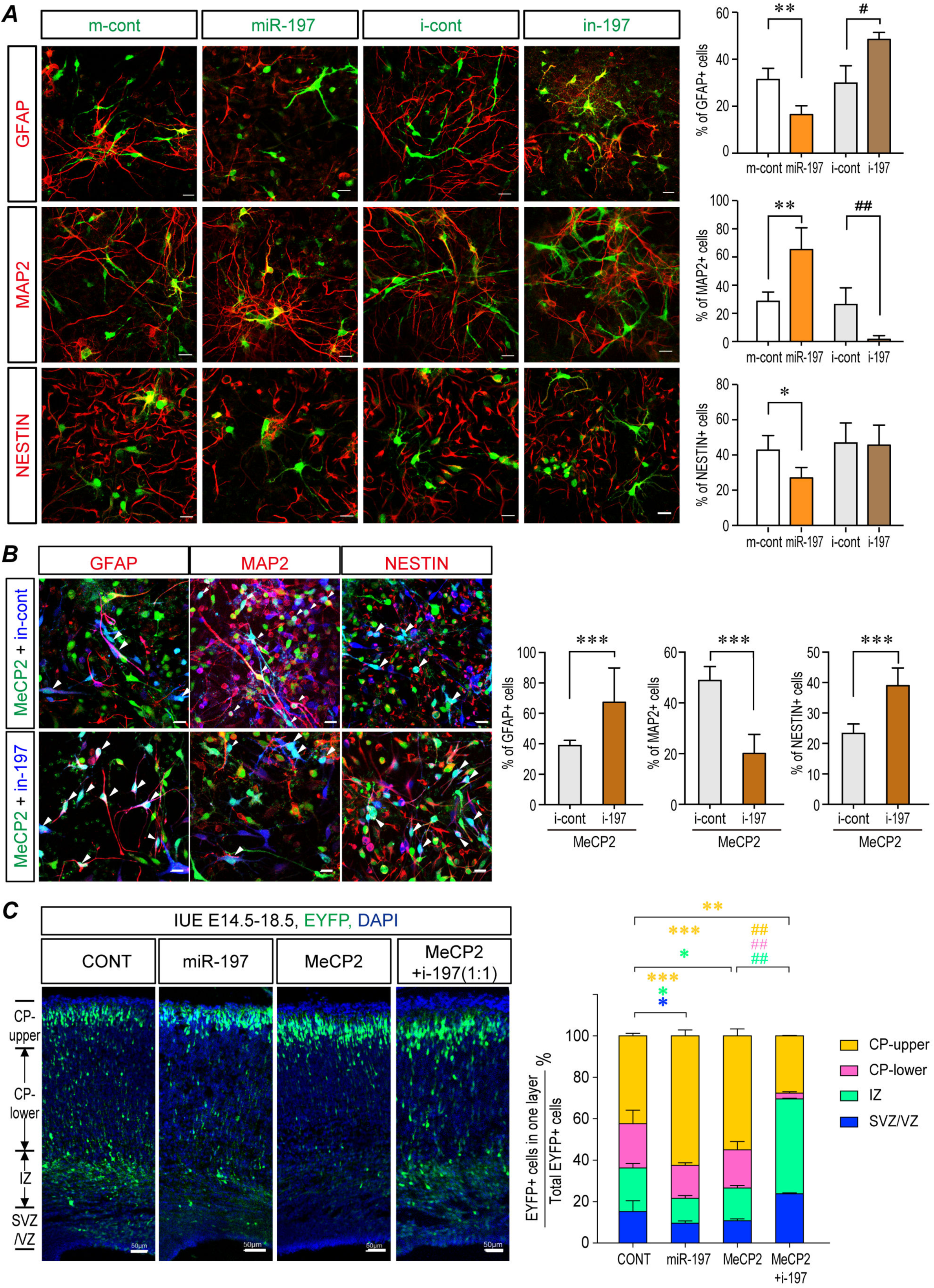
miR-197 promotes neurogenesis and the inhibitor of miR-197 reverses the effect of MeCP2 overexpression. **(A)** Primary NPCs from C57BL/6 mouse were infected with assorted lentivirus to over-express either miR-197, miR-control, miR-197 inhibitor (i-197), or inhibitor-control. 72 hrs later, cells were subjected for immunofluorescent staining. The infected cells are labeled with pseudo-colored green. The staining for MAP2, GFAP, and NESTIN are labeled with pseudo-colored red. The percentage of Nestin^+^, MAP2^+^ and GFAP^+^ cells within infected cells are shown on the right side respectively in. N≥4. **(B)** Primary mouse NPCs were infected with assorted lentivirus to overexpress MeCP2 with either i-197 or i-cont. 72 hrs later, cells were subjected for immunofluorescent staining for MAP2, GFAP, and NESTIN. The MeCP2 and control viruses contain EGFP and showed here as green. The i-197/i-cont viruses infected cells are pseudo-color labeled as blue. The staining for MAP2, GFAP, and NESTIN are labeled with pseudo-colored red. The percentage of Nestin^+^, MAP2^+^ and GFAP^+^ cells within green and blue double positive cells (cyan in nucleus, white arrow head) are shown on the right side, respectively. N=9. **(C)** Overexpression of miR-197 mimicked the effect of MeCP2 and miR-197 inhibitors (i-197) blocked the effect of overexpressed MeCP2 in WT C57BL/6 mouse fetal brain by IUE experiments. N=9. Representative brain sections are presented on the left side, and the statistical analyses for cells in each layer on the right panel. All statistical data are presented as means ± SEM. * or # *p*<0.05, ** or ## *p*<0.01, *** or ### *p*<0.001. Scale bar is 50μm.

Similar results were observed in the mouse fetal brain. The effect of overexpressing miR-197 alone was almost identical to MeCP2 overexpression in WT C57BL/6 mouse cortex, with more EYFP^+^ cells reached the upper CP layer and less EYFP^+^ cells stayed in the VZ/SVZ layer than in the controls (Fig. 5C). More importantly, i-197 blocked the effect caused by MeCP2 overexpression in WT brain. That is, the number of EYFP^+^ cells reaching the upper CP layer was reduced by i-197 (Fig. 5C). Expression of i-197 in Tg(*MECP2*) mice could also reverse the effect of elevated MeCP2 expression, as significantly more EYFP^+^ cells remained in the VZ/SVZ and IZ layers and fewer EYFP^+^ cells reached the upper CP layer (Fig. S5B). These data strongly suggest that miR-197 plays a key role in MeCP2 mediated neurogenesis during cortical development.

### *MECP2* mutations identified in a Chinese ASD cohort failed to promote neurogenesis

A targeted-sequencing of *MECP2* gene exons was performed in a Han Chinese cohort consisting of 288 ASD patients and 369 controls. Five rare missense *MECP2* mutations were identified in six male ASD patients that were not observed among any of our controls (Table 1). These mutations were c.590C>T, c.695G>C, c.1112A>G, c.1180G>A, and c.1282G>A, which correlated to T197M, G232A, H371R, E394K and G428S in the human MeCP2-e1 protein (mouse MeCP2-e2) (Fig. S6), represents the isoform highly expressed in the brain (32, 49). The patients carrying these mutations showed a spectrum of symptoms, and the patient carrying the G428S mutation presented the most severe symptoms including severe intellectual disability with no functional language skills, and other medical conditions such as sleep disturbances (Table 1). According to ExAC, three C-terminal mutations (H371R, E394K and G428S) were not previously identified in the East Asian population (Total 4327 samples, with Male/Female=2,016/2,311) (39). The recurrent mutation H371R in our cohort represents a novel mutation in all ExAC populations, a database which currently contains sixty thousand individuals (39). Therefore, we focused on those C-terminal mutations in the follow up studies.

**Table 1.**
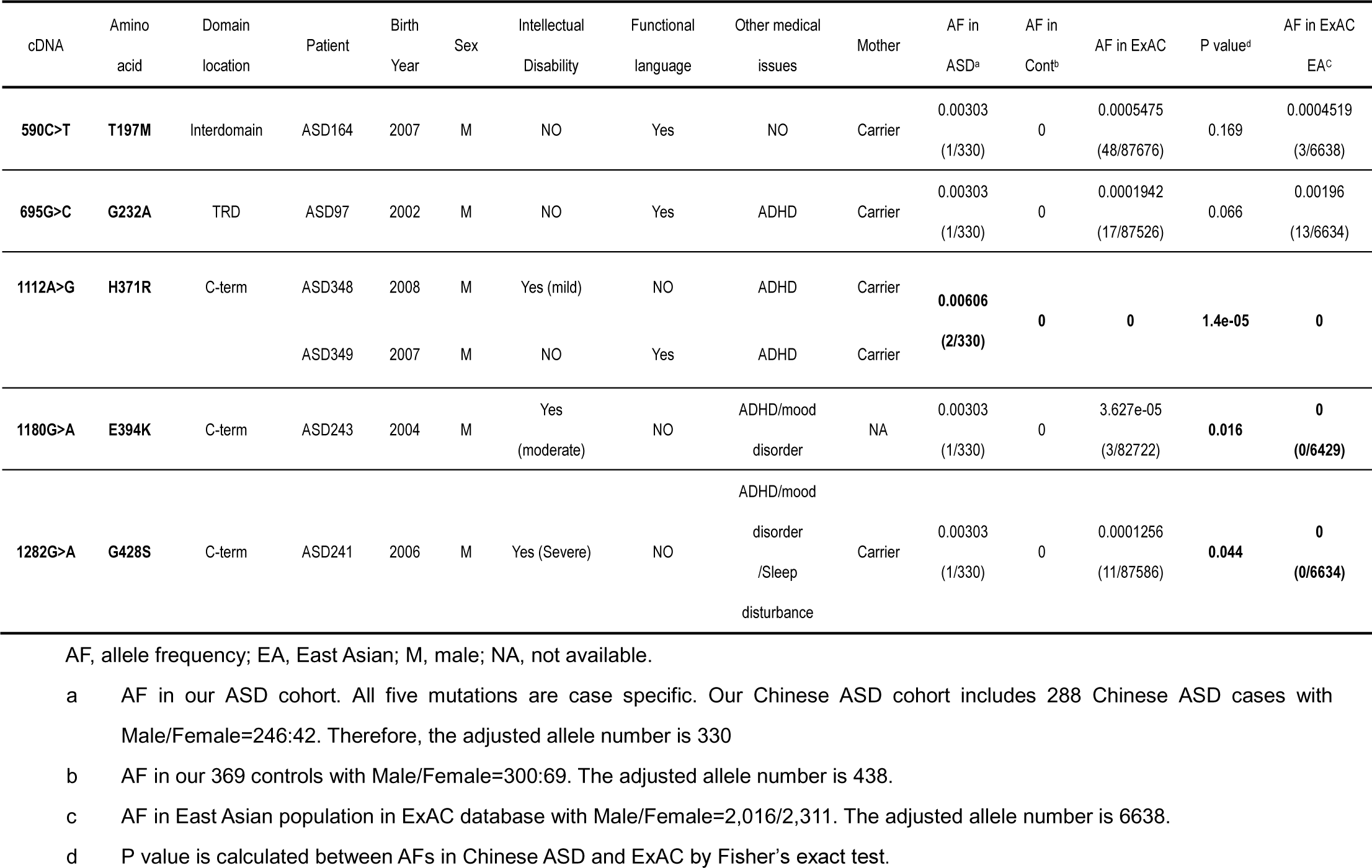
Rare Mutations of *MECP2* detected in Chinese ASD cohort.

We investigated the effect of three C-terminal mutants on NPCs differentiation. Although these mutations did not affect the expression levels of MeCP2 when overexpressed in NPCs (Fig. S7), they failed to promote neurogenesis in both NPCs and mouse embryonic cortex (Fig. 6B&C). Both MeCP2^H371R^ and MeCP2^E394K^ showed loss-of-function effects similar to that of MeCP2^380X^, a previously reported loss-of-function truncation mutant (20) in IUE experiments, as they resulted in fewer EYFP^+^ cells reaching the upper CP layer, and more EYFP^+^ cells remaining in the VZ/SVZ layer compared to the WT MeCP2 electroporated brain (Fig. 6B). To be noticed, MeCP2^G428S^ produced a more severe phenotype when overexpressed in mouse embryonic cortex with most of the MeCP2^G428S^ overexpressing cells remaining within the VZ/SVZ layers in the mouse cerebral cortex, and very few cells reached the CP at E18.5 (Fig. 6B).

**Figure 6.**
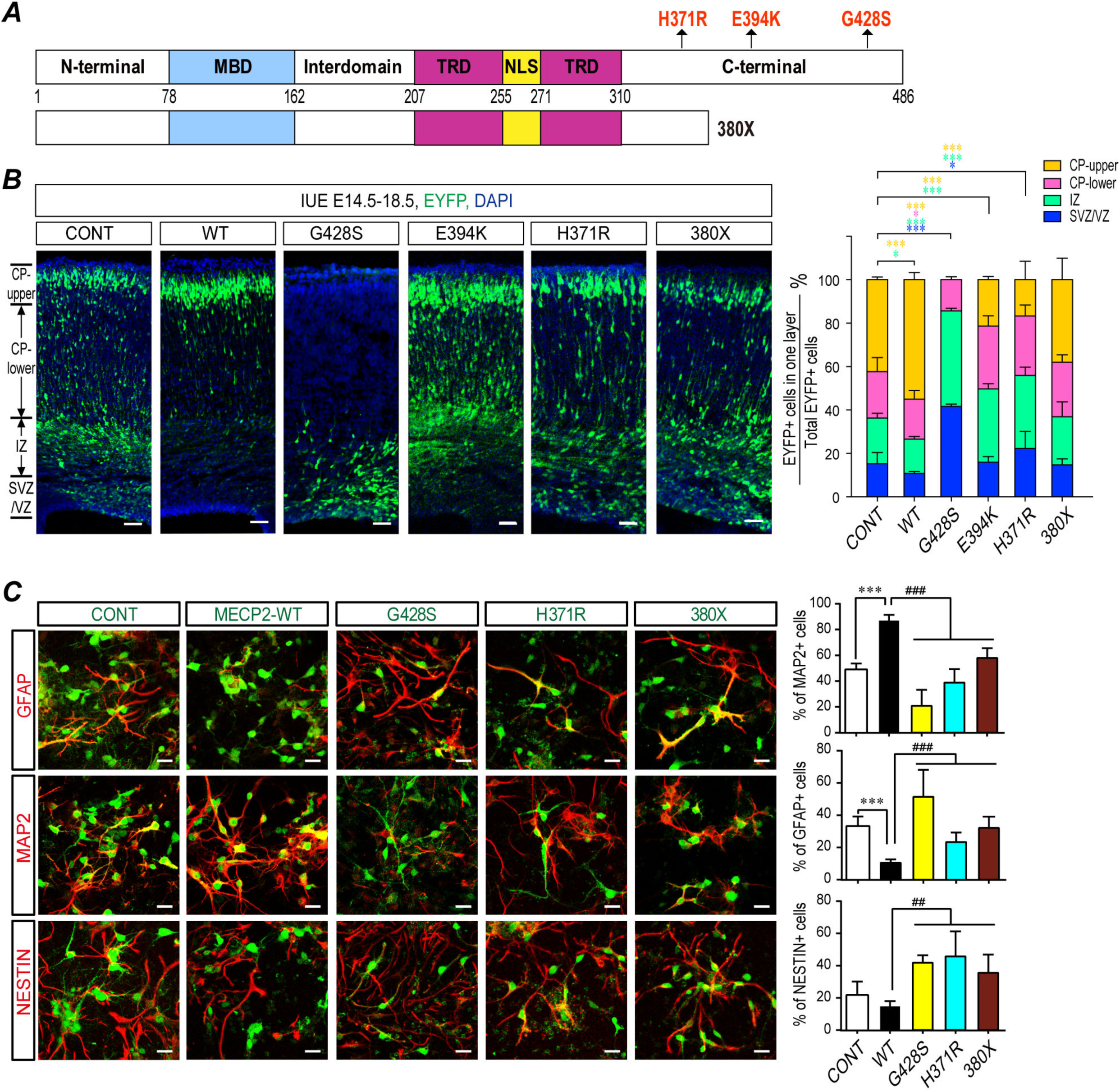
ASD-related MeCP2 mutants lose the effect on promoting neurogenesis. **(A)** A scheme illustration for positions of the ASD-related *MECP2* mutations, H371R, E394K, G428S, and a reported truncated mutant 380X. **(B)** Fetal WT C57BL/6 mouse brains were electroporated at E14.5 to overexpress either WT MeCP2 or ASD-related MeCP2 mutants. Samples were collected at E18.5 for sectioning and immunostaining. Each condition was repeated in four different embryos from four pregnant dams (N=4). Representative brain sections are presented on the left and the statistical analyses for cells in each layer are shown on the right. **(C)** Primary mouse NPCs were infected with assorted lentivirus to overexpress WT MeCP2, MeCP2^H371R^, MeCP2^G428S^, and MeCP2^380X^. 72 hrs later, cells were subjected for immunofluorescent staining for MAP2, GFAP, and NESTIN. The infected cells are labeled with EGFP (green). The staining for MAP2, GFAP, and NESTIN are labeled with pseudo-colored red. The percentage of MAP2^+^, GFAP^+^, and Nestin^+^ cells within EGFP+ cells are shown on the right, respectively. N=9. All statistical data are presented as means ± SEM. * or # *p*<0.05, ** or ## *p*<0.01, *** or ### *p*<0.001. Scale bar is 50μm.

Since the patient with the MeCP2^G428S^ mutation presented the most severe symptoms compared to other two mutations, and the MeCP2^G428S^ mutant had the strongest effect in IUE experiment, we went on to inspect the effects of MeCP2^G428S^ mutation in the *in vitro* NPC differentiation assay. Overexpression of the MeCP2^G428S^ mutant led to a significantly reduced percentage of MAP2^+^ cells (~21%) and a higher percentage of Nestin^+^ cells (~42%), compared to either WT MeCP2 (~87% MAP2^+^ and ~15% Nestin^+^ cells, respectively) or the control empty vectors (~49% MAP2^+^ and ~22% Nestin^+^ cells, respectively) (Fig. 6C). The MeCP2^G428S^ mutant demonstrated a gliogenesis promoting effect during NPC differentiation as indicated by a higher percentage of GFAP^+^ cells (~51%) than WT MeCP2 (~11%), or the control (~33%) (Fig. 6C).

### MiR-197 reversed the neurogenesis defects caused by ASD-related MeCP2 mutants

We also investigated the effect of those MeCP2 mutants on the levels of miR-197 and ADAM10. The results showed that three MeCP2 C-terminal mutants lost the ability to up-regulate pre-miR-197 and matured miR-197 (Fig. 7A&B), or to down-regulate ADAM10 and NICD expression (Fig. S8). Again, MeCP2^G428S^ showed some gain-of-function effect evidenced as a significant up-regulation in NICD level in both mouse NPCs and human U251 cells (Fig. S8). However, when miR-197 was co-transfected with those three MeCP2 mutants, it was able to down-regulate the expression of ADAM10 even in presence of the MeCP2 mutants (Fig. 7C). Finally, when miR-197 was co-electroporated with MeCP2G428S into E14.5 WT mouse brains, the abnormal distribution of cells detected in MeCP2^G428S^ expressing cortex at E18.5 could be almost completely reversed by exogenous miR-197 (Fig. 7D).

**Figure 7.**
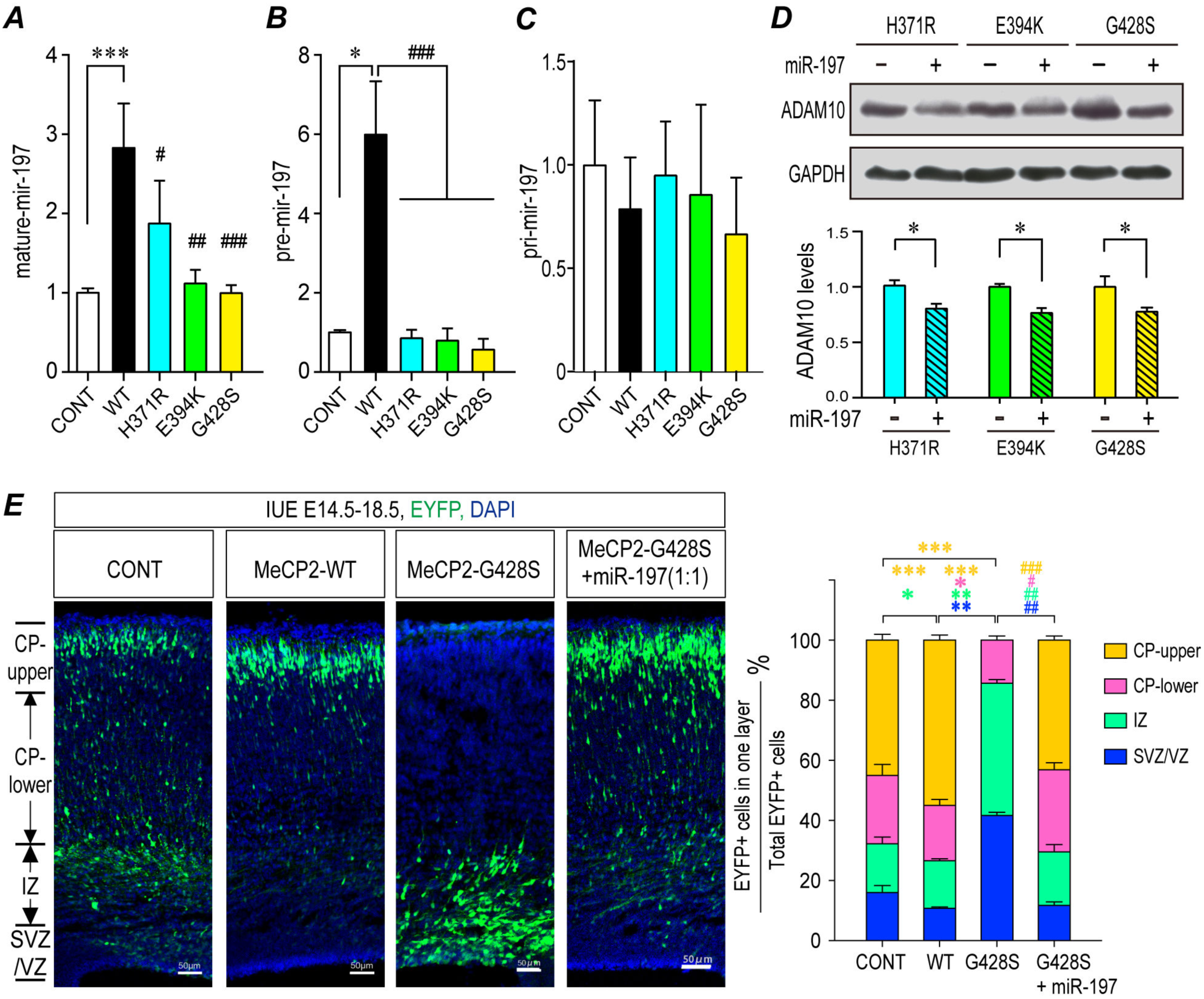
MiR-197 reverses the neurogenesis defects induced by ASD-related MeCP2 mutants. U251 cells were transfected with either WT or mutant MeCP2 expressing plasmids and cultured for 24hrs. RNA from these cells was subjected to qPCR for either mature miR-197 **(A)**, pre-miR-197 **(B)**, or pri-miR-197 **(C)**. N=6 for mature miR-197, N=4 for pre and pri-miR-197. **(D)** U251 cells were transfected with mutant MeCP2 expressing plasmids and either control or miR-197 plasmid. Cells were cultured for 24hrs and cell lysates were subjected to western blot for ADAM10. N≥5. Representative blots are shown on the left side and statistical analyses are shown on the right side, respectively. **(E)** Overexpression of miR-197 reversed the differentiation defects caused by MeCP2^G428S^ in electroporated fetal brain. Each condition was repeated in four different embryos from four pregnant dams (N=4). Representative brain sections are presented on the left, and the statistical analyses for cells in each layer is shown on the right. All statistical data are presented as means ± SEM. * or # *p*<0.05, ** or ## *p*<0.01, *** or ### *p*<0.001. Scale bar is 50μm.

## Discussion

Duplication or mutations of the *MECP2* gene has been shown to cause MDS and RTT, respectively. 100% of MDS patients and >60% of RTT patients express autistic behaviors, yet the underlying mechanism remains elusive. It has been proposed that altered NSC/NPC differentiation may contribute to the etiology of ASD (50). Again, the mechanism by which MeCP2 affects NSC/NPC differentiation is unknown. In this study, we demonstrated that the dosage and function of MeCP2 is tightly related to embryonic NSC/NPC differentiation. We showed in a MDS animal model that elevated MeCP2 expression promotes neurogenesis by down-regulating ADAM10 and suppressing NOTCH signaling. Moreover, we identified three rare missense mutations in *MECP2* in an ASD cohort and found that they are loss-of-function mutants involved in regulating NSC/NPC differentiation. Our results revealed that the defects in a regulatory axis involving MeCP2, miR-197, ADAM10, and NOTCH signaling is associated with the etiology of MDS and/or ASD. Not only can ADAM10 and/or the miR-197 inhibitor inhibit the neurogenesis promoting effect of elevated MeCP2 levels in the Tg(*MECP2*) mouse brain, but additionally miR-197 can reverse the differentiation defects caused by the ASD-related MeCP2 mutants both *in vitro* and *in vivo*. These results indicate that miR-197 must be a critical molecule for the etiology of MDS and ASD.

The role of MeCP2 in NSC/NPC cell fate determination is complex and varies by its dosage, spatial, and temporal pattern. For example, there are studies reporting that *Mecp2* deficiency does not affect NPCs fate in the *Mecp2-/y* mouse embryonic cortex (51, 52), rather it affects the neuron dendritic arborization and complexity in the mouse brain (51-53). However, overexpression of MeCP2 not only promotes neurogenesis of NPCs isolated from embryonic cortex (42), but also causes premature neurogenesis in the developing chick neural tube (33). Herein, we also observed enhanced neurogenesis in Tg(*MECP2*) mouse and in WT embryonic cortex with ectopic expression of MeCP2. Taken together, *MECP2* duplication probably has more severe effects during the early embryonic stages. These observations are consistent with the clinical features of MDS patients, who have severe phenotypes starting from very early infantile stages (9, 10).

A surprising finding is that miR-197, which we identified as a crucial molecule downstream of MeCP2, does not regulate human *ADAM10*-3’UTR through the conventional miRNA seed sequence. Rather, miR-197 binds to an unconventional site on *ADAM10*-3’UTR. The fact that this unconventional binding site but not the predicted miR-197 seed sequence is conserved between human and mouse suggests that the interaction between miR-197 to *ADAM10*-3’UTR is more similar to a siRNA. Although the mouse miR-197 was removed from miRBase in 2014 based on the fact that “the sequence does not map in a stem-loop region of the genomic sequence or any known mouse transcript sequence”; however, we noticed that small RNA libraries were used to identify miR-197 in both human and mouse in the original paper by Landgraf *et al.* (54). Indeed, a small RNA could be PCR amplified from E18.5 mouse brain by using has-miR-197 specific primers (Fig. S4C&D). It is possible that there might be a small RNA with a similar sequence and function that exists in the mouse brain, even though it does not fit with the typical stem-loop structure of miRNAs.

Another interesting observation is that pre-miR-197 is up-regulated by MeCP2. Previous studies have shown that MeCP2 can both down-regulate and up-regulate miRNAs (20, 21, 55). Down-regulation on miRNA, such as pre-miR-134, is due to MeCP2’s binding to DGCR8 to interfere with the assembly of Drosha/DGCR8 miRNA processing complex (20). Indeed, pre-miR-134 is still down-regulated by MeCP2 in our system (Fig. 3L). Therefore, the MeCP2 regulatory mechanisms on miR-197 and miR-134 probably are unique to each miRNA. Recently, Tsujimura *et al.* found that miR-199a is up-regulated by MeCP2 through binding to Drosha and DDX5 (21). It is of interest that both pri-miR-199a (21) and pri-miR-197, but not pri-miR-134, can be experimentally pulled down by MeCP2 (Fig. S9). It is possible that when MeCP2 binds to a pri-miRNA, it facilitate the miRNA processing, but when MeCP2 cannot bind to a pri-miRNA, it interferes miRNA processing. Future studies to investigate whether other miRNAs regulated by MeCP2 also fit such rules are critically important.

It has been proposed that rare inherited mutations are important in the etiology of ASD (56-60). Indeed, most of the ASD cases with the *MECP2* mutations in our cohort inherited the mutation from their asymptomatic carrier mother (Table 1). As all those ASD patients are boys, mutated *MECP2* on the X chromosome would affect them more severely than their mothers. Although we cannot conclude that those mutations are causative without further genetic analysis of their whole genome, it is noticeable that the H371R mutation has two recurrent cases in our ASD patients (2/288), but does not exists in 61,075 controls from both ExAC and our cohort. Taken together with the fact that H371R has similar *in vivo* and *in vitro* effects as does the truncation mutation 380X, it is highly possible that H371R is a loss-of-function mutation with a high likelihood of being causative for ASD. The G428S mutation showed more severe effects on NPCs differentiation when compared to the 380X mutation. How this mutation changes MeCP2’s function will need further investigation. One possibility is the G428 to S mutation might induce a new phosphorylation site on MeCP2, as previous studies have shown that the phosphorylation status of MeCP2 affects its functions (28, 61, 62). There are 4 hemizygotes (male carriers) for G428S in the ExAC database from South Asian and Latino populations. However, the different genetic background of each population could affect the prevalence and susceptibility of any given mutation. If we look at only the East Asian population in ExAC, all three C-terminal mutations are novel out of a total of 4696 East Asian genomes. Whole-exome sequencing or even whole genome sequencing of our ASD cohort in combination with other ASD sequencing data sets are likely to be needed in the future to fully understand the genetic importance of those *MECP2* mutations.

In conclusion, we uncovered a novel mechanism by which MeCP2 regulates NSC/NPC differentiation through the NOTCH pathway via miR-197. Together with a recent whole-exome sequencing study of over 1000 ASD cases (63) and our observations, deregulation of the NOTCH pathway could be important to the etiology of MDS and ASD.

## Material & Methods

### Animal Housing and Genotyping

Both C57BL/6 mice and FVB-Tg(*MECP2*)1Hzo/J mice were maintained in the animal facility at the Institute of Developmental Biology & Molecular Medicine, Fudan University. The protocol was approved by the Committee on the Ethics of Animal Experiments of Fudan University. The genotyping of FVB-Tg(*MECP2*)1Hzo/J mice were determined by PCR following the protocols from Jackson Laboratory website.

### Analyses of mouse cortical neurogenesis *in vivo*

For quantification of cell fate in WT and FVB-Tg(*MECP2*)1Hzo/J mice at E18.5 and P7, the regions of the primary somatosensory cortex were identified and the numbers of Sox+, Tbr2+, Satb2+ or Tbr1+ cells were counted in each vertical column with 100 μm width. All quantifications were performed with 4 brain sections from at least 3 animals. Data are presented as the mean ± SEM and statistical significance was assessed using unpaired Student’s t test.

### Primary mouse NPCs isolation and differentiation assay

NPCs were isolated from E12.5 C57BL/6 or FVB mouse embryonic cortex. NPCs isolation and culture methods are based on previous reports (42, 64). NPCs were either transfected with X-tremeGENE HP DNA transfection reagent (Roche, 06366236001) or infected with different lentivirus (Obio Technology (Shanghai) Corp., Ltd.). 72 hours post-transfection or infection, NPCs cells were either lysed for western blot or subjected to immunofluorescent staining and imaging with a Zeiss LSM700 microscope. All films from western blot were scanned and analyzed with Quantity ONE based on intensity or were directly measured with Tanon gel image software. The results were normalized to its corresponding loading control GAPDH.

### *In utero* electroporation and cell counting

*In utero* electroporation was performed as previously described (65, 66). About 1.5 μl of DNA mix was injected into each embryo. The ratio of either expressing plasmid or empty vector to pEYFP was 6:1. The ratio of (miR-197 inhibitor or ADAM10): (WT MeCP2): pEYFP was 3:3:1. The ratio of miR-197, MeCP2^G428S^, and pEYFP was 3:3:1. After electroporation and recovery, E18.5 embryos were collected and sectioned at 20μm and processed for further immunofluorescent analyses. Nuclear cell staining with DAPI was used to define different sub-regions of the cerebral cortex based on cell density, as previously described (65). The percentage of EYFP^+^ cells in each layer was calculated based on total number of EYFP^+^ cells in the same brain section. At least three sections from one brain were collected and at least four different embryos obtained from 3-4 different pregnant dams were collected for each group for statistical analyses..

### Plasmids and Lentivirus

Human *MECP2*-e1 cDNA was purchased from Origene. Point mutations were generated by mutagenesis PCR. The lentivrus for human WT MeCP2, H371K, G428S, 380X, sh-MeCP2 and sh-control were all purchased from Obio Technology (Shanghai) Corp., Ltd. The corresponding mutations were also constructed into rat *MECP2*-e2 cDNA with a HA tag in pRK5 vector and used in IUE.

Two different regions of human *ADAM10* 3’-UTR were cloned into XbaI/FseI sites of pGL3 plasmid to generate luciferase reporter constructs. A10-I-WT contains 1461-1765 of the *ADAM10* 3’-UTR, which covers the predicted miR-197 binding site (1568-1574) by TargetscanHuman (http://www.targetscan.org/vert_71/) (47). A10-II-WT contains 4-405 of human *ADAM10* 3’-UTR, which covers the predicted binding site (186-201) by RNAhybrid analysis (48).

### qRT-PCR

Total RNA from NPCs or U251 was extracted with RNAeasy kit (QIAGEN, Cat# 74104). The levels of ADAM10 mRNA were determined using the qRT-PCR kit (Takara RR036A and RR820A) with *GAPDH* as internal control. The levels of mature miR-197 were determined with All-in-One^™^ miRNA qRT-PCR Detection Kit (GeneCopoeia, QP015). The levels of pri-miR-197 were detected with TaqMan pri-microRNA assay kit (Applied Biosystems) following reverse transcript with Toyobo ReverTra Ace-α kit (Toyobo). The levels of pre-miR-197 were detected with miScript Precusor Assay (QIAGEN) following reverse transcript with miScript II RT Kit (QIAGEN).

### RNA Immunoprecipitation

U251 cells were transfected with empty vector, WT MeCP2, or ASD-related MeCP2 mutant expressing plasmids. 24 hours later, cells were collected for cross-linking, lysing, and sonication and were subjected to the RNA immunoprecipitation assay using previously described protocols (21, 67). Anti-MeCP2 antibody (ab2828) and rabbit immunoglobulin G (IgG) antibodies were used. The immunoprecipitated RNA was analyzed by qRT-PCR described before.

### Biotinylated Micro-RNA Pull Down Assay for Identifying miRNA Targets

Biotinylated double stranded miRNA-197 and its scrambled control miRNA (B02003) (both Biotin-labeled at 3’ end) were purchased from GenePharma. U251 cells were transfected with control miRNA and miR-197 at a final concentration of 100 nM with RNAimax (Invitrogen^TM^, 13778150). 24 hours post transfection, whole cell lysates were harvested and subjected to RNP pull down followed with RT-PCR. The miRNA enrichment was calculated as follow: Bi-miR-197 pull-down for ADAM10 3’UTR /Scramble control pull-down for ADAM10 3’UTR = X, Bi-miR-197 input/Bio-Scramble control input = Y, Fold binding = X/Y. At least three independent experiments with a minimum of three replicates each time were performed for each set.

### Statistical analyses

All experiments were repeated at least three times and the statistical significance was evaluated. Data are expressed as mean ± SEM. Statistical differences were calculated by two-tailed unpaired t test for two datasets and ANOVA followed by Bonferroni post hoc test for multiple datasets using Prism (GraphPad Inc., La Jolla, CA). *p*<0.05 was considered statistically significant.

### Human subjects

Blood samples from 288 ASD patients (mean age 6.1±3.1 years, 85% male) were collected between 2007 and 2010 at the Shanghai Mental Health Center of Shanghai Jiaotong University. Protocols were reviewed and approved by the Ethics Committee of Fudan University and Shanghai Jiaotong University prior to the commencement of the study. Written informed consent from the parents or guardians of the children was obtained prior to inclusion in the study. The 369 controls (mean age 19 years, 81% male) were unrelated healthy volunteers from the freshman student class at Fudan University (2010), which were ethnically and gender-matched from the same geographical area. DNA was extracted and all samples were pooled together for targeted sequencing on the 2kb upstream, 5’-UTR, 3’-UTR, and coding regions of *MECP2* gene was carried out at GBP Biotechnology (Jiangsu, China). Sanger sequencing was performed to confirm those 5 missense mutations in the 6 ASD cases (Fig. S8A).

## Acknowledgement

We would like to thank Dr. Xiongli Yang, Dr. Xiang Yu, and Dr. Xiaohui Wu for their valuable suggestions on our manuscript.

This work was supported by the following grants to Drs. Hongyan Wang and Yufang Zheng: the National Key Basic Research Program of China (2016YFC1000502), the NSFC (81430005, 31771669, 81741048, 31521003), the National Basic Research Program (973) of China (2013CB945404), the Commission for Science and Technology of Shanghai Municipality (17JC1400902 and 14JC1401000), the 863 program of China (2014AA021104); and the Changjiang Scholarship to Dr. Richard H. Finnell.

The authors have declared that no conflict of interest exists.

## Supplementary material and methods

### FVB-Tg(MECP2)1Hzo/J mice Genotyping

The genotyping of FVB-Tg(*MECP2*)1Hzo/J mice were determined by PCR following the protocols from Jackson Laboratory website: https://www2.jax.org/protocolsdb/f?p=116:5:0::NO:5:P5_MASTER_PROTOCOL_ID,P5_JRS_CODE:14245,008679.

### MeCP2 Plasmids and Lentivirus

Human *MECP2*-e1 cDNA was purchased from Origene (RC202382). Each point mutant was generated by mutagenesis PCR. The lentivrus for human WT MeCP2 (HQ664), H371K (H4971), G428S (H4972), 380X (H4973), sh-MeCP2 (HQ363) and sh-control (H101) were all purchased from Obio Technology (Shanghai) Corp., Ltd.

### miRNA inhibitors and mimics

Plasmids for hsa-miR-197 precursor (HmiR0013-MR04), hsa-miR-197 inhibitor (HmiR-AN0287-AM01), and scrambled negative control clone (CmiR0001-MR04) were purchased from GeneCopoeia, Inc. USA. The lentivirus for miR-197 (H4976) and miR-control (H32), i-197 (Y3496) and i-control (Y008) were all purchased from Obio Technology (Shanghai) Corp., Ltd. MiR-197 mimics (B01001) and scrambles (B04001); MiRNA inhibitors (B03001) and their negative controls (B04003) were purchased from Shanghai GenePharma Co,Ltd..

### Antibodies

Antibodies used in this study were as follow: from Abcam: ADAM10 (ab1997), ADAM17 (ab13535), NICD (ab8925), GFAP (ab7260), NOTCH1 (ab27526), DLL-1 (ab76655), JAG-1 (ab7771), Nestin (ab6142), MAP2 (ab11268, ab32454), GFAP (ab7260), NeuN (ab177487), Tuj1 (class III beta tubulin, ab78078), Satb2 (ab92446), SOX2 (ab97959), TBR1 (ab31940), TBR2 (ab23345), HA (ab9110); and Alexa Fluor^®^ 647-labeled goat anti-Rabbit (ab150079), Alexa Fluor^®^ 647-labeled goat anti-mouse (ab150115), or Alexa Fluor^®^ 488-labeled goat anti-Rabbit (ab150077), Alexa Fluor^®^ 488-labeled goat anti-mouse (ab150117). Anti-MeCP2 and GAPDH antibody was purchased from Cell Signaling (CST, #3456 & #2118S). All HRP-conjugated secondary antibodies were purchased from KangChen Bio-tech Inc., Shanghai, China.

### Primers

The primers for hsa-mir-197-3p (HmiRQP0287) and the internal control snRNA U6 gene (HmiRQP9001 for human U6 and MmiRQP9002 for mouse U6) were also purchased from GeneCopoeia Inc. Primers for pri-miR-197 (4427012), pri-miR-134 (4427012), and the internal control GAPDH gene (4331182) were purchased from Applied Biosystems. Primers for pre-miR-197 (Hs_mir-197_PR_1 miScript Precursor Assay, MP00001302), pre-miR-134 (Hs_mir-134_PR_1 miScript Precursor Assay, MP00000847) and the internal control U6 gene (Hs_RNU6-2-11 miScript Primer Assay, MS00033740) were purchased from QIAGEN.

Point mutations in MECP2 were generated with the following primers.

H371R: 5’-GAGCACCACCACCATCACCGCCACTCAGAGTCCCCAAAG-3’,

5’-CTTTGGGGACTCTGAGTGGCGGTGATGGTGGTGGTGCTC-3’;
E394K: 5’-CCACCTGAGCCCAAGAGCTCCGAGG-3’

5’-CCTCGGAGCTCTTGGGCTCAGGTGG-3’;
G428S: 5’-CACTGGAGAGCGACAGCTGCCCCAAGGAGCC-3’

5’-GGCTCCTTGGGGCAGCTGTCGCTCTCCAGTG-3’

The primers used for PCR cloning A10-II-WT are as follow.

F: 5’-CAGTCTAGACAGCTTTTGCCTTGGTTCTT-3’;
R: 5’-ACTGGCCGGCCGGTCGAGCCTCCTAGCCTTGATTG-3’.

The primers used to generate A10-II-Mut are as follow:

F1: 5’-CAGTCTAGACAGCTTTTGCCTTGGTTCTT-3’;
R1: 5’-AAGAAAATTGGGTTCCTTTTAATTGGAATTTTCAGGCTTT-3’;
F2: 5’-AAAGCCTGAAAATTCCAATTAAAAGGAACCCAATTTTCTT-3’;
R2: 5’-ACTGGCCGGCCGGTCGAGCCTCCTAGCCTGATTG-3’.

The primers used for qRT-PCR were:

ADAM10: 5’-CCTGCCATTTCACTCTGTCATTTA-3’

5’-GTGCCCGGGCTCCTTCCTCTACTC-3’;
GAPDH: 5’-ACAGCAACTCCCACTCTTCCACCT-3’
  5’-TTGCTCAGTGTCCTTGCTGGGG-3’.

The primers used for RT-PCR ADAM10 3’UTR in Biotinylated Micro-RNA Pull Down Assay were:

5’-CAGTCTAGACAGCTTTTGCCTTGGTTCTT-3’;
5’-ACTGGCCGGCCGGTCGAGCCTCCTAGCC TTGATTG-3’

### The inhibitors of miRNA

hsa-miR-197-3p, 5’-GCUGGGUGGAGAAGGUGGUGAA-3’;
hsa-miR-221-3p, 5’-GAAACCCAGCAGACAAUGUAGCU-3’;
hsa-miR-199a-5p, 5’-GAACAGGUAGUCUGAACACUGGG-3’,
hsa-miR-222-3p, 5’-ACCCAGUAGCCAGAUGUAGCU-3’,
hsa-miR-137, 5’-CUACGCGUAUUCUUAAGCAAUAA-3’,
hsa-miR-193a-3p, 5’-ACUGGGACUUUGUAGGCCAGUU-3’,
hsa-miR-184, 5’-ACCCUUAUCAGUUCUCCGUCCA-3’,
hsa-miR-187-3p, 5’-CCGGCUGCAACACAAGACACGA-3’.

### Biotinylated Micro-RNA Pull Down Assay for Identifying miRNA Targets

Biotinylated double stranded miRNA-197 and its scrambled control miRNA (B02003) (both Biotin-labeled at 3’ end) were purchased from GenePharma. U251 cells were transfected with control miRNA and miR-197 at a final concentration of 100 nM with RNAimax (Invitrogen^TM^, 13778150). 24 hours post transfection, whole cell lysates were harvested in the lysis buffer (Sigma-Aldrich, I8896) supplemented with protease inhibitor (Thermo Scientific^TM^, 78430) and RNase inhibitor (Thermo Scientific^TM^, EO0381). After centrifugation, the supernatants were collected. 50 μl Streptavidin-Dyna beads (Dyna beads M-280 streptavidin) (InvitrogenTM, catalog#: 11205D) and 10 μl yeast tRNA (AmbionTM, AM7119) were added to 500 μl cell lysates and incubated at 4 °C overnight. The next day, 750 μl Trizol and 250 μl water were added per tube and kept at −20°C overnight. The mixture was thawed the following day and 200 μl chloroform was added and vortexed for 45 sec. The mixture was centrifuged at 4°C 18,000 × g for 15 min. The upper layer was transferred to new tubes to which equal volumes of 2-propanol and 5 μl glycoblue (Invitrogen^TM^, AM9515) were added. This was incubated at RT for 10 min and then centrifuged at 4°C 18,000 × g for 15 min. The supernatant was very carefully discarded and the pellet was washed twice with 1 ml pre-chilled 70% ethanol. The pellet was air dried and resuspended in 20 μl nuclease free water. RT-PCR: 20 μl RNA from biotin-labeled sample and 1 μg RNA from the input sample was used for mRNA RT reactions. The miRNA enrichment calculation: Bi-miR-197 pull-down for ADAM10 3’UTR /Scramble control pull-down for ADAM10 3’UTR = X, Bi-miR-197 input/Bio-Scramble control input = Y, Fold binding = X/Y. At least three independent experiments with a minimum of three replicates each time were performed for each set.

---

**Figure S1.**
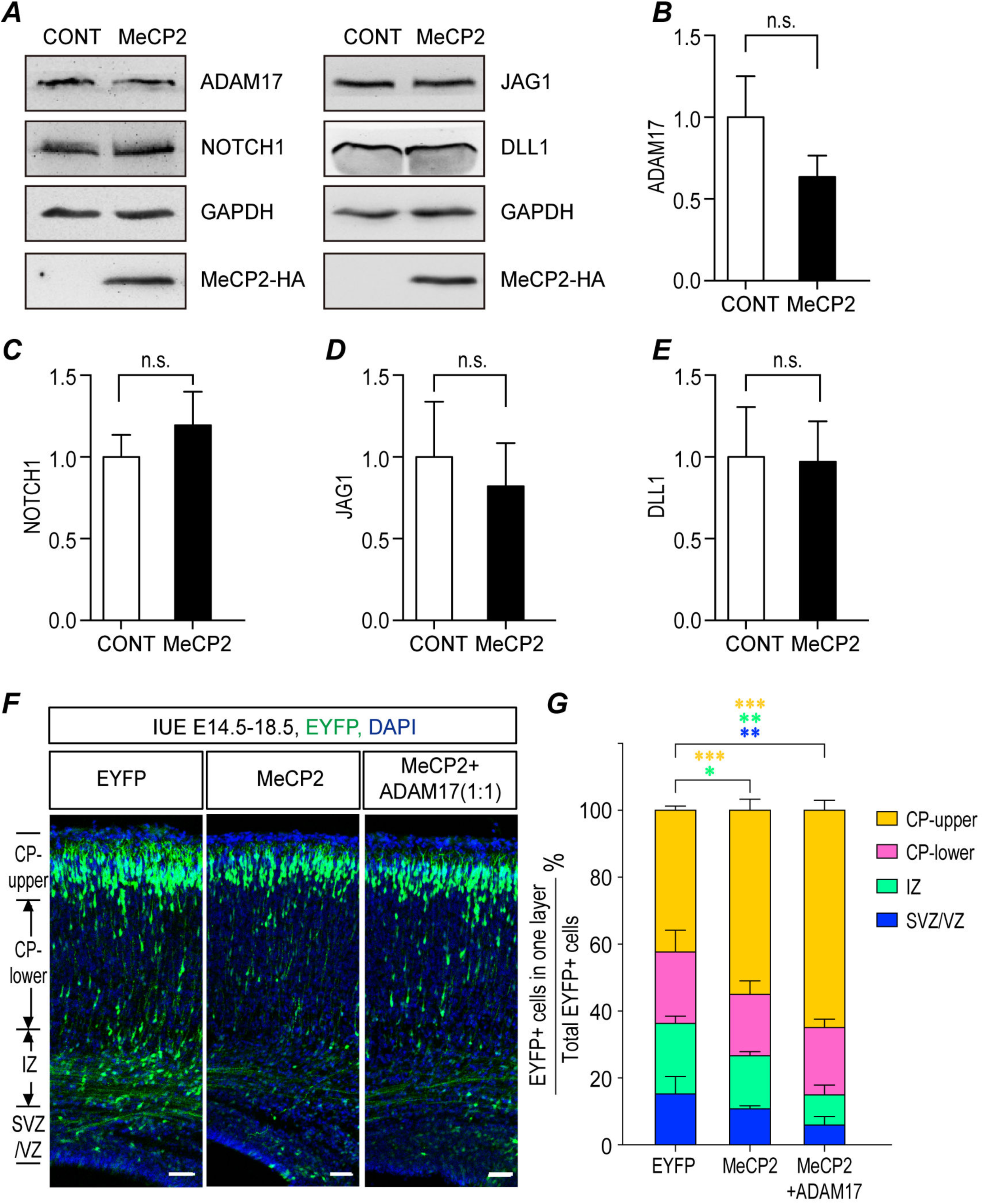
ADAM17 could not reverse MeCP2 promoted neurogenesis. **(A-E)** Mouse primary NPCs isolated from E12.5 mouse embryonic cortex were transfected with either empty vector or MeCP2 expressing plasmids and cultured for 72hrs. Cell lysates were subjected to western blot analysis for other components of NOTCH pathway except ADAM10 and NICD. N≥5. Representative blots are shown in **(A)** and statistical analyses for each component was presented as follow, **(B)** ADAM17, **(C)** NOTCH1, **(D)** JAG1, and **(E)** DLL1. **(F-G)** Overexpression of ADAM17 could not reverse the differentiation effects of MeCP2 in electroporated embryonic brain. Fetal mouse brains were electroporated at E14.5 and the samples were collected at E18.5 for sectioning and immunostaining **(F)**. Each condition was repeated in four different embryos from four pregnant dams (N=4). Representative brain sections are presented on the left side, and statistical analyses for cells in each layers are shown in **(G)**. All statistical data represent means ± SEM. ** *p*<0.01, *** *p*<0.001. Scale bar is 50μm.

**Figure S2.**
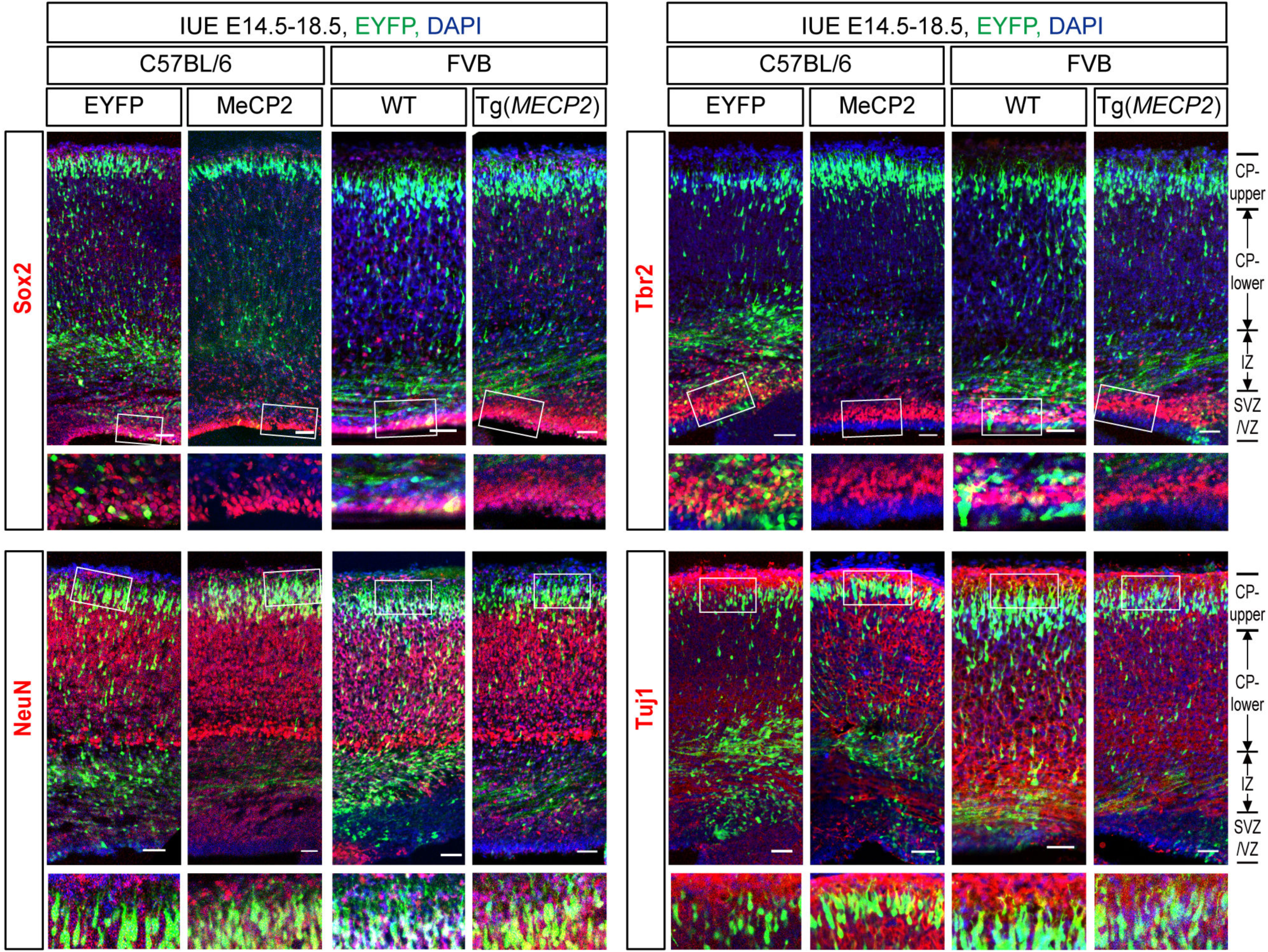
MeCP2 overexpression promotes neurogenesis in IUE brains. C57BL/6 or FVB fetal mouse brains were electroporated at E14.5 and the samples were collected at E18.5 for sectioning and immunofluorescent staining with progenitor markers Sox2 and Tbr2, and neuronal markers NeuN and Tuj1 (red). DAPI (blue) was used for nuclear staining. C57BL/6 brains were electroporated with EYFP and MeCP2 expressing plasmids or control vector. FVB brains were electroporated with EYFP alone. Each condition was repeated in four different embryos from four pregnant dams (N=4). Representative sections were presented. Enlarged images for white boxes are presented on the bottom. Scale bar is 50μm.

**Figure S3.**
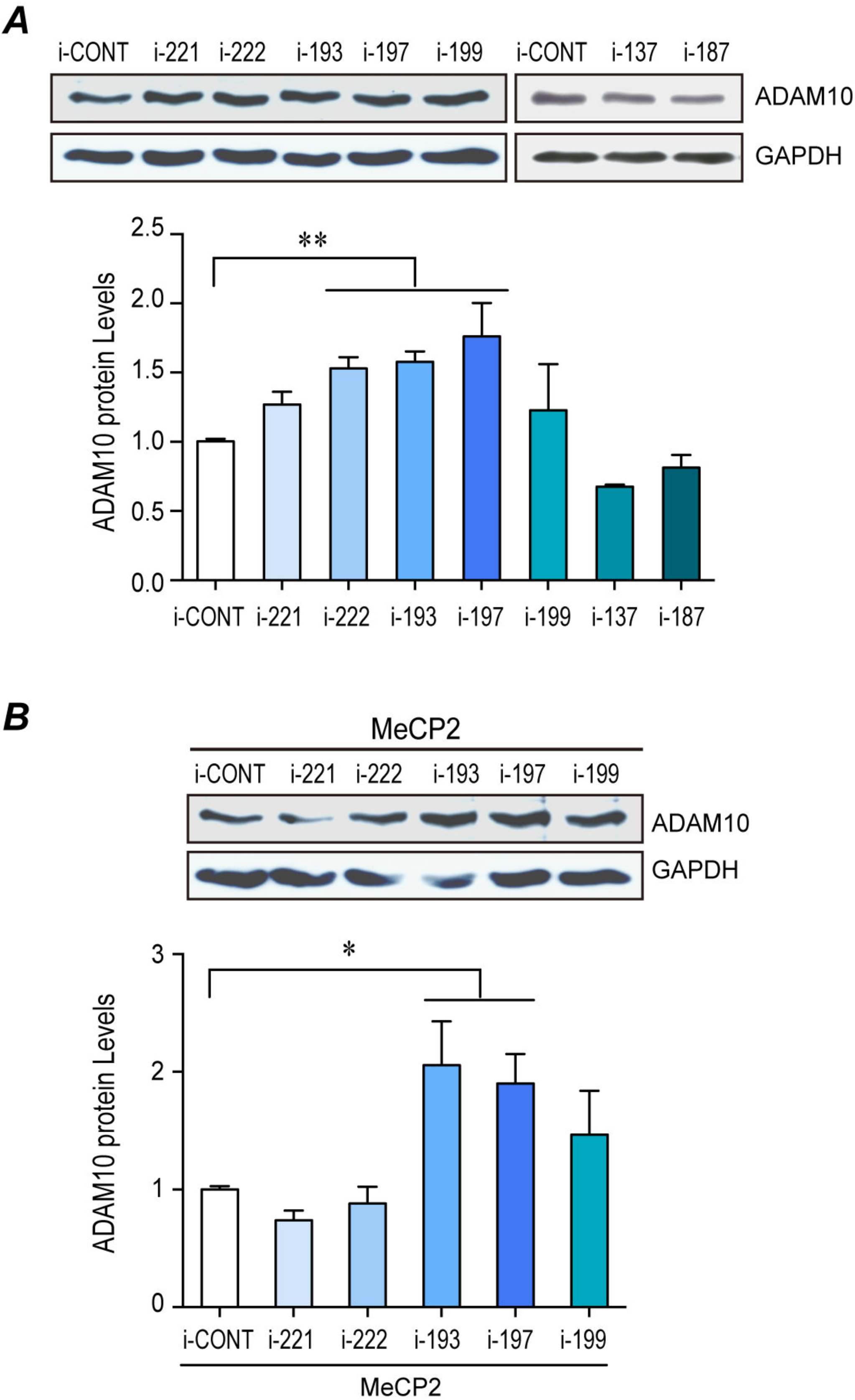
MiR-197 inhibitor reversed the down-regulation effect of ADAM10 by MeCP2. **(A)** Inhibitors for miR-221, miR-222, miR-193, miR-197, miR-199, miR-137, and miR-187 were transfected into U251 cells and the levels of ADAM10 protein were examined. **(B)** Inhibitors for miR-221, miR-222, miR-193, miR-197, and miR-199 were co-transfected with WT MeCP2 into U251 cells, and the levels of ADAM10 protein were examined. Representative blots are shown on the top panels, and statistical analyses for ADAM10 protein are shown on the lower panels. All data represent means ± SEM. N≥3, * *p*<0.05, ** *p*<0.01.

**Figure S4.**
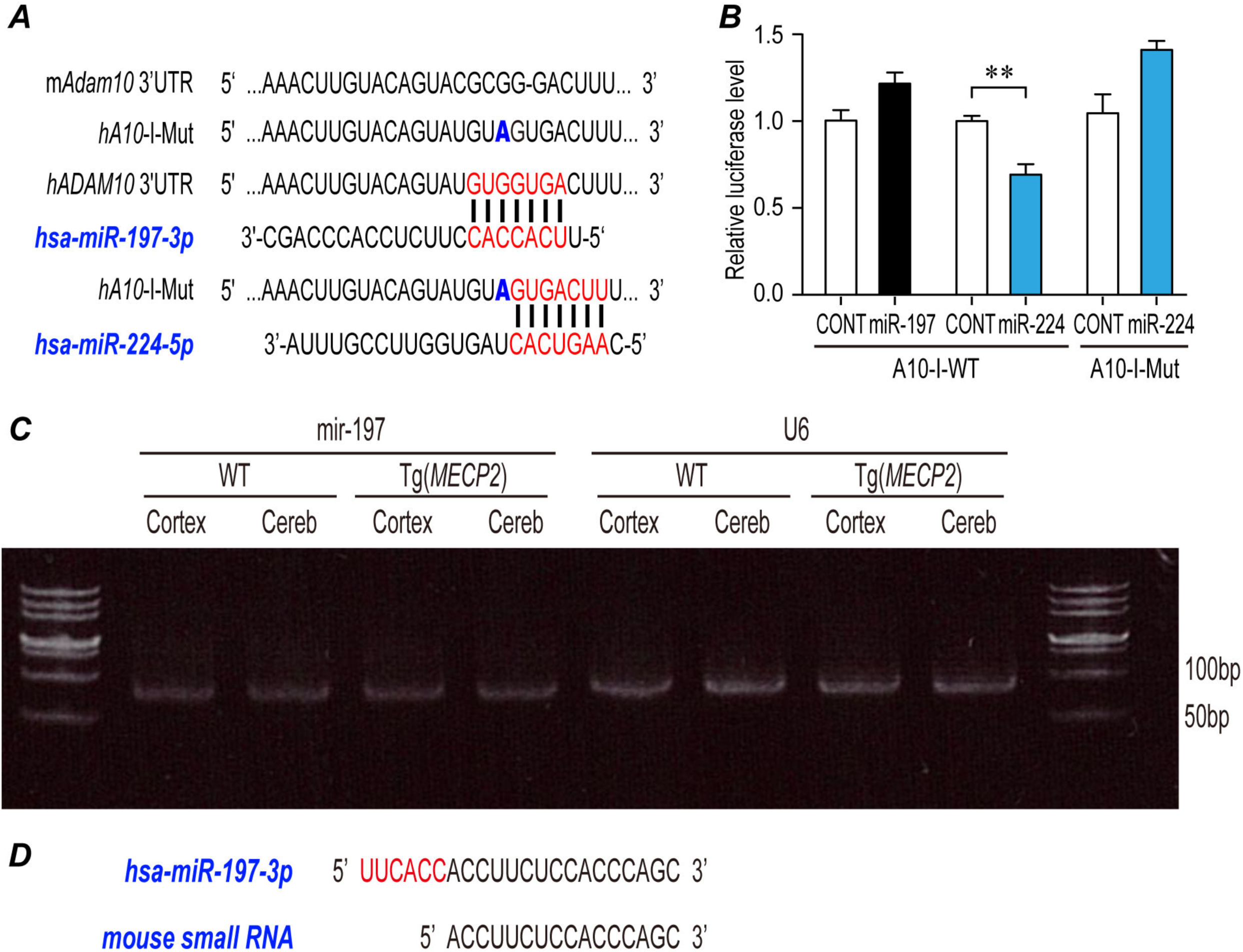
A miR-197 like small RNA could be RT-PCR amplified from E18.5 mouse cortex. **(A)** A 7mer-m8 miR-197 binding site at position 1568-1574 of human *ADAM10* 3’-UTR was predicted by TargetscanHuman, which is poorly conserved between human and mouse. The alignment of miR-197 to human *ADAM10* 3’-UTR and miR-224 to A10-I-Mut were illustrated. **(B)** Luciferase reporter assay showed that miR-224 but not miR-197 could down-regulate the expression of A10-I-WT. The point mutation A10-I-Mut is not sensitive to miR-224 anymore. All data represent means ± SEM. N≥3, ** *p*<0.01. **(C)** Fetal cortex and cerebellum from E18.5 WT and Tg(*MECP2*) FVB mice were dissected out. RNA was extracted and subjected to RT-PCR with either hsa-miR-197 specific primer or control Rnu6 (U6 small nuclear RNA) primers. 8μl products and the DNA ladder DL500 (Takara) were run on 12% PAGE gel without urea and stained with Gel-Red for 40min. **(D)** The PCR product was cloned into T-vector and sent for sequencing. The small RNA has 16 identical nucleotides to the 3’ side of has-miR-197.

**Figure S5.**
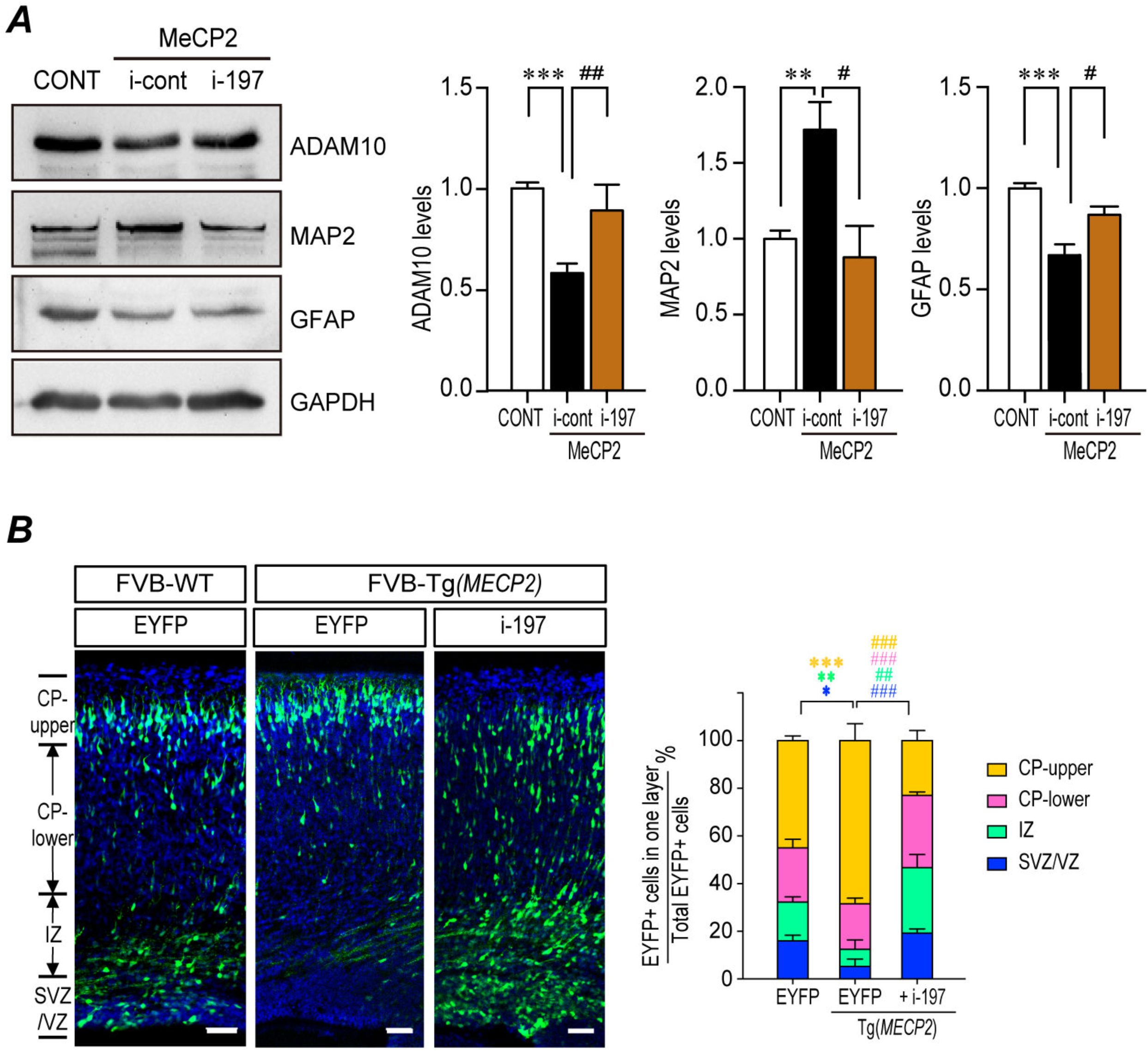
The miR-197 inhibitor reverses the effect of elevated MeCP2 in mouse brain. **(A)** Primary NPCs from C57BL/6 mouse were transfected with MeCP2 expressing plasmid together with either inhibitor control or i-197. Cell lysates were subjected to Western blot analysis for MAP2, GFAP, and ADAM10. N≥4. Representative blots are shown on the left side and statistical analyses are shown on the right side, respectively. **(B)** The neurogenesis stimulated in the Tg(*MECP2*) mouse fetal brain could be blocked by overexpressing miR-197 inhibitors (i-197). Fetal mouse brains were electroporated at E14.5 and collected at E18.5 for sectioning and immunostaining. N=4. DAPI (blue) was used for nuclear staining. Representative staining images were presented. Scale bar is 50μm. All statistical data represent means ± SEM. *, # *p*<0.05, **, ## *p*<0.01, ***, ## *p*<0.001.

**Figure S6.**
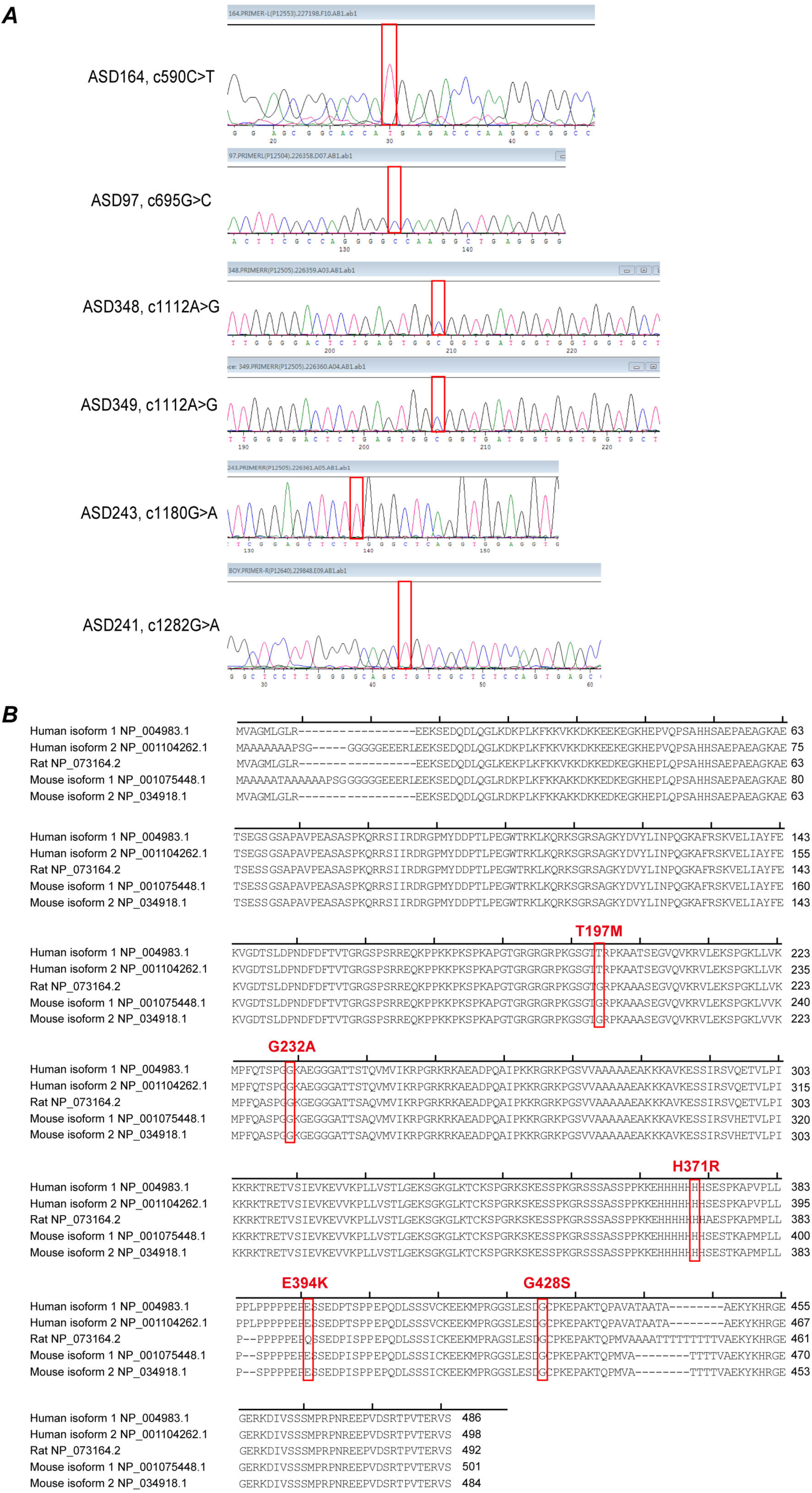
The sequencing data and alignment of autism-related MECP2 mutations. **(A)** The MECP2 mutations in the ASD patients were confirmed by Sanger sequencing. **(B)** MeCP2 is a highly conserved protein. Sequence alignment of human MeCP2 e1 and e2, mouse MeCP2 e1 and e2, and rat MeCP2 e2 proteins by NCBI online alignment software. The mutated amino acids are highlighted in red box.

**Figure S7.**
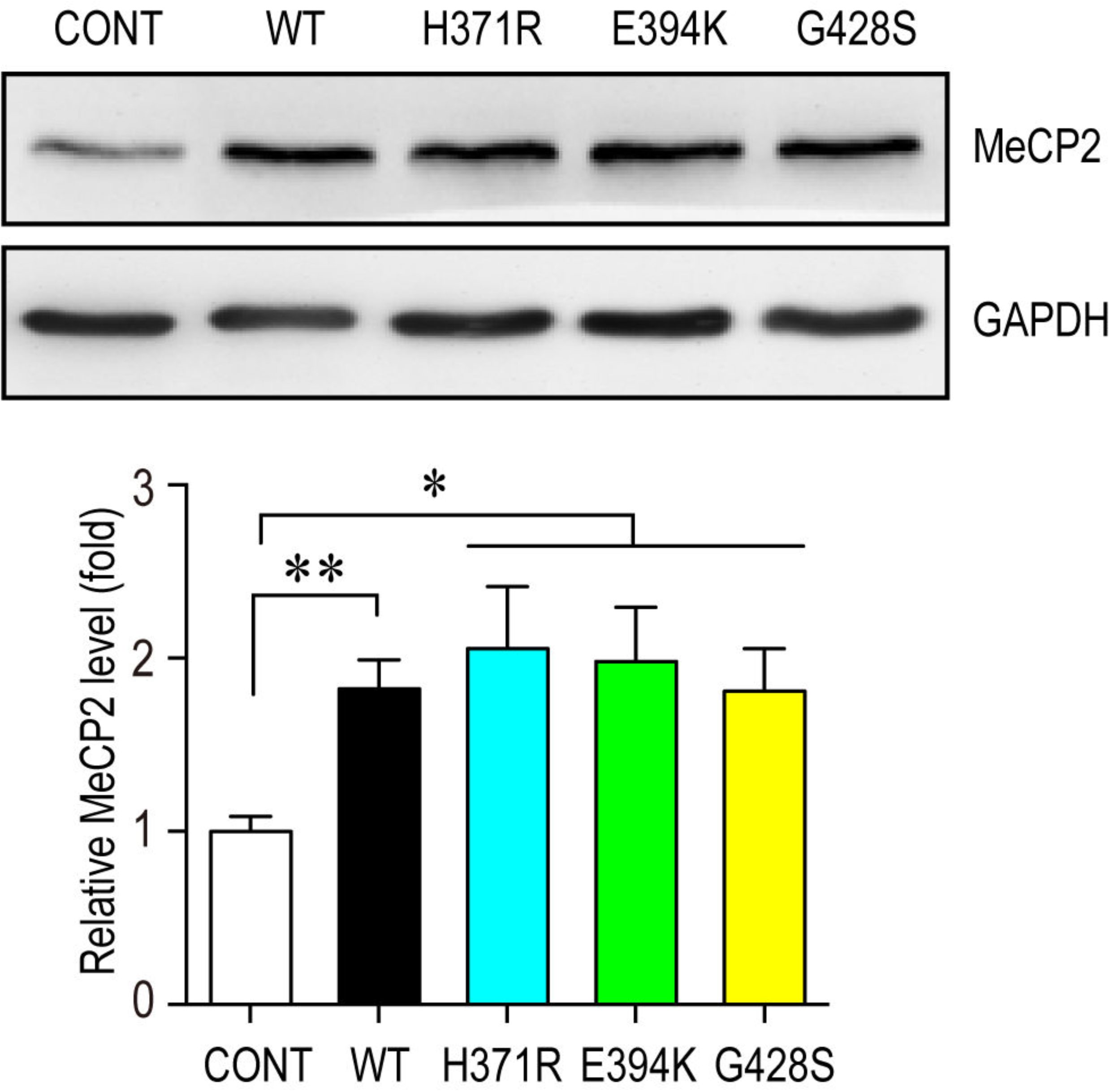
Exogenous expressing of MeCP2 mutants in NPCs were similar to WT-MeCP2. Mouse primary NPCs isolated from C57BL/6 mouse E12.5 embryonic cortex were transfected with either control vector or different MeCP2 expressing plasmids and cultured for 72hrs. Cell lysates were subjected to western blot analysis for MeCP2 protein level and GAPDH was used as a loading control. Representative blot is shown on the top panel, and statistical analysis for MeCP2 levels are shown in the bottom panel. N=3. All statistic data represent means ± SEM. * *p*<0.05, ** *p*<0.01.

**Figure S8.**
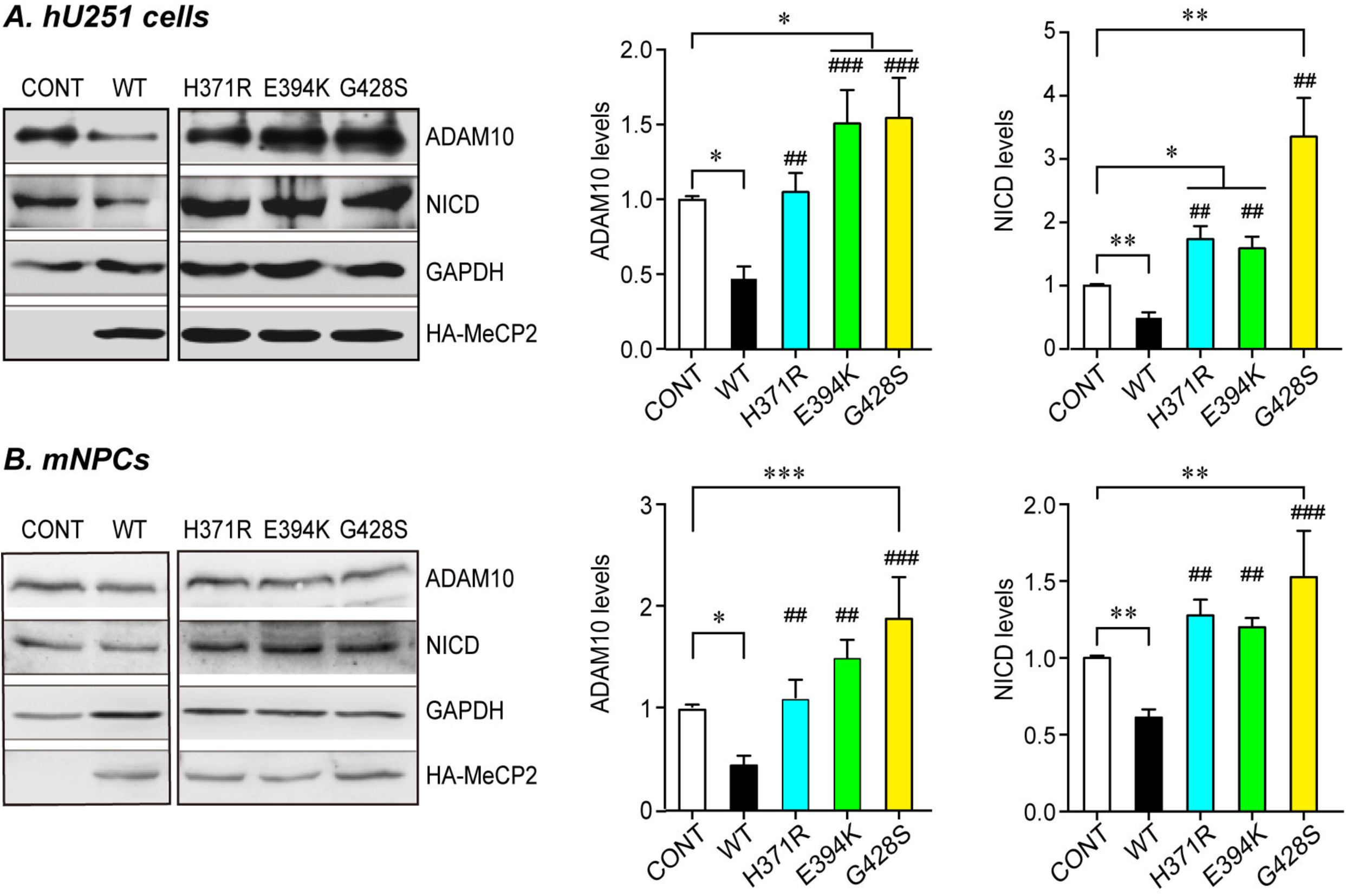
Three C-terminal mutant MeCP2 failed to down-regulate ADAM10 and NICD. Human U251 cells (**A**) or mouse primary NPCs (**B**) were transfected with either WT or three C-terminal MeCP2 mutant expressing plasmids and cultured for 24hrs or 72hrs, respectively. Cell lysates were subjected to western blot analysis for ADAM10 and NICD. Representative blots are presented on the left side and statistical analyses for cells in each layer are shown on the right side. GAPDH was used as internal control. All data represent means ± SEM. Effects of the mutations are either compared to control (indicated with *), or WT MeCP2 (indicated with #). N≥4, All statistical data represent means ± SEM. * or # *p*<0.05, ** or ## *p*<0.01, *** or ### *p*<0.001.

**Figure S9.**
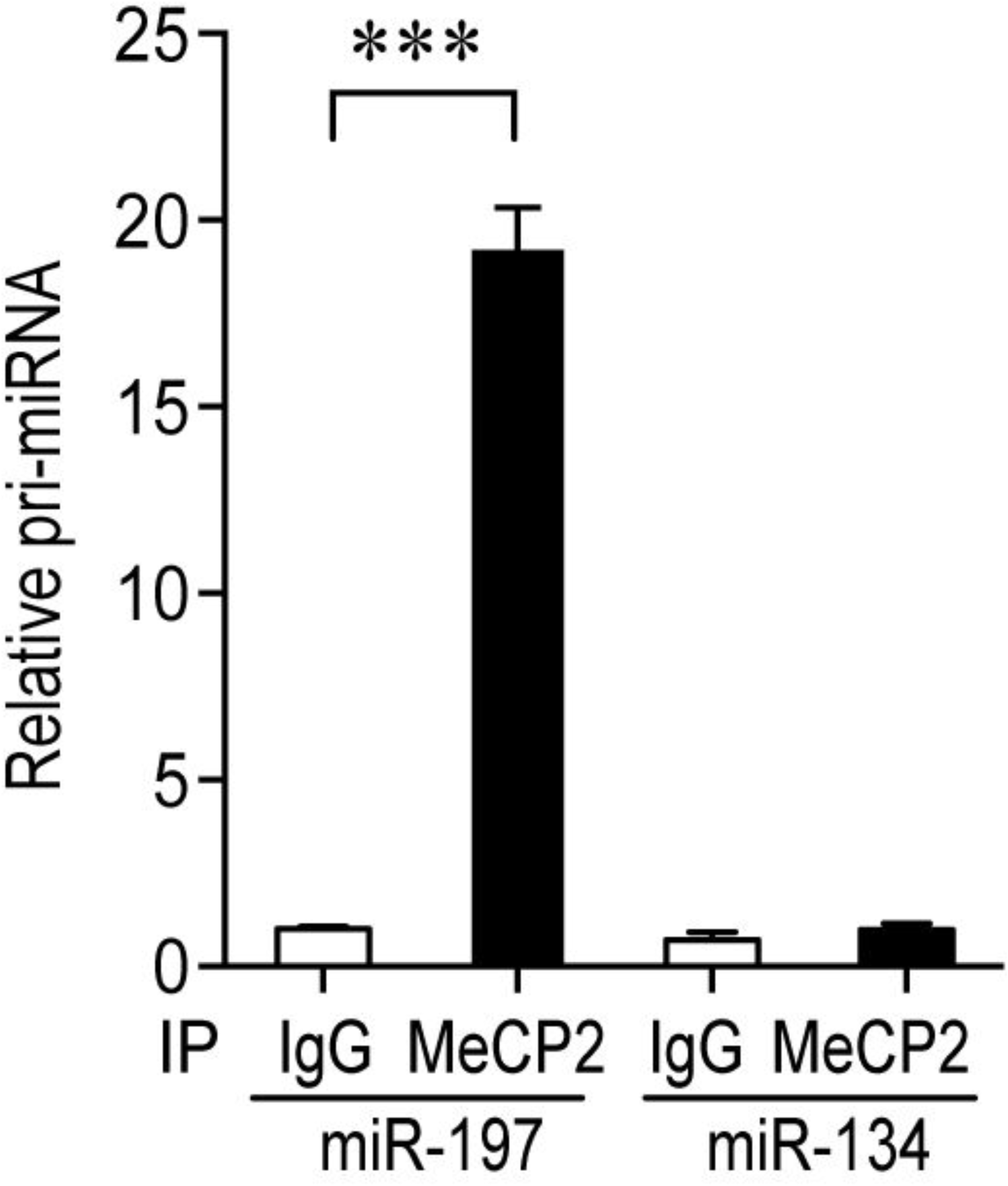
MeCP2 interacts to miR-197 but not miR-134 and MeCP2^G428S^ does not bind to miR-197 anymore. U251 cells transfected with MeCP2 expressing plasmids were also subjected to RNA-IP with MeCP2 antibody or control IgG. The levels of pri-miR-197 and pri-miR-134 co-precipitated with MeCP2 were quantified by qRT-PCR. N≥5, all statistical data represent means ± SEM. *** *p*<0.001.

